# A method for massively scalable inference of phylogenetic networks

**DOI:** 10.1101/2025.05.05.652278

**Authors:** Nathan Kolbow, Sungsik Kong, Claudia Solís-Lemus

## Abstract

Recent advances in sequencing technologies have enabled large-scale phylogenomic analyses. While these analyses often rely on phylogenetic trees, increasing evidence suggests that non-treelike evolutionary events, such as hybridization and horizontal gene transfer, are prevalent in the evolutionary histories of many species, in which case tree-based models are insufficient. Phylogenetic networks can capture such complex evolutionary histories, but current methods for accurately inferring them lack scalability. Implicit network inference methods are fast but lack biological interpretability. Here, we introduce a novel method called InPhyNet that merges a set of non-overlapping, independently inferred level-1 networks into a unified topology, achieving linear scalability while maintaining high accuracy under the multispecies network coalescent model. We prove that a statistically consistent and scalable pipeline for inferring phylogenetic networks can be constructed with InPhyNet. Using simulation, we infer networks with up to 200 taxa and show that divide-and-conquer pipelines utilizing InPhyNet allow for accurate network inference at scales and speeds previously unseen. Re-analyzing a phylogeny of 1,158 land plants with InPhyNet, we recover well-accepted reticulate events and illustrate how InPhyNet enables large-scale analyses of biologically meaningful reticulate phylogenies at previously unprecedented scales.

## Introduction

Advances in sequencing technologies have increased the availability of biological genomic data for phylogenetic analyses. These analyses are foundational to our understandings of many biological processes such as cancerous tumor progression (Schwartz and Schäffer, 2017), infectious disease spread (Ciccozzi et al., 2019), drug discovery (Searls, 2003), conservation biology (Véron et al., 2019), and crop domestication in agriculture (Sang and Ge, 2007). To date, the backbone of these analyses has been phylogenetic trees: bifurcating tree models that represent evolutionary relationships based on two descendant species sharing a single common ancestor. However, growing bodies of evidence suggest that some evolutionary events, such as hybridization and horizontal gene transfer, are prevalent in the histories of many species (Boto, 2010). These phenomena diverge from the strictly bifurcating structure enforced by phylogenetic trees, rendering such trees inadequate for modeling them.

Phylogenetic networks can accurately model non-treelike, reticulate evolution, but are much more computationally intensive to infer due to their increased model complexity, higher dimensional parameter space, and a larger set of possible topologies. Indeed, without restrictions, the space of network topologies is infinite. State-of-the-art maximum composite likelihood (sometimes referred to as pseudo-likelihood) inference methods such as Species Network applying Quartets (SNaQ) (Solís-Lemus and Ané, 2016), Phylogenetic Network Estimation using SiTe patterns (PhyNEST) (Kong et al., 2025), or PhyloNet’s maximum pseudolikelihood (MPL) option (Yu and Nakhleh, 2015) offer reasonable speeds and high accuracies but can only handle up to approximately 30 taxa. More recent research to increase this upper bound has been fruitful, leading to methods like FastNet (Hejase et al., 2018) or PhyloNet’s divide-and-conquer option (Zhu et al., 2019), the latter of which can handle up to 81 taxa. However, these methods still have the fundamental issue of high-degree polynomial runtime complexity.

This runtime issue can be circumvented by inferring implicit networks, a category of networks that does not directly model biological phenomena and instead summarizes conflicting signals in data. Methods for inferring implicit networks such as NeighborNet (Bryant and Huson, 2023) or Consensus Networks (Holland et al., 2004) are often employed when researchers wish to analyze hundreds or thousands of taxa with a network model (Forster et al., 2020). However, such analyses are often inappropriate given their lack of biological grounding (Kong et al., 2016; Sánchez-Pacheco et al., 2020).

Here, we present InPhyNet, a novel method for constructing arbitrarily large semi-directed phylogenetic networks that scales *linearly* with respect to the number of taxa in the network. InPhyNet works by merging a set of level-1, semi-directed, binary phylogenetic networks with the guidance of a pairwise dissimilarity matrix. This results in a semi-directed, binary, tree-based phylogenetic network that is not necessarily level-1. This approach facilitates the accurate, efficient inference of phylogenetic networks with over 1,000 species while leveraging highly accurate maximum likelihood (ML) and maximum composite likelihood (MCL) methods.

We show that such a pipeline can be statistically consistent for the underlying species network under the multispecies network coalescent model (Yu et al., 2014). The practicality of this theory is verified with a rigorous simulation study that evaluates InPhyNet’s performance under realistically complex conditions.

Lastly, we re-evaluate the phylogeny of 1,158 land plant species (One Thousand Plant Transcriptomes Initiative, 2019) by inferring reticulations in portions of the phylogeny that were poorly resolved in their estimated phylogenetic tree. We find support for previously well-accepted instances of reticulate evolution as well as evidence for a controversial placement of Gnetales, a division of plants, within gymnosperms. Our method is available as an open-source Julia package at https://github.com/NathanKolbow/InPhyNet.jl.

All scripts and results used in the simulations presented here are available at github.com/NathanKolbow/InPhyNetSimulations, and all original data are deposited in the Dryad Digital Repository, available at https://doi.org/10.5061/dryad.h44j0zq0b.

## Materials AND Methods

### Scalable Inference Framework

This framework aims to make phylogenetic network inference feasible for larger numbers of input taxa. To achieve this, we use a divide-and-conquer strategy: the original, intractable problem is divided into smaller, tractable subproblems, and a novel method is then applied to merge the solutions of these subproblems into a final result. For a set of input data *G* of arbitrary type on the set of taxa *X*, the framework works as follows:

1. Decompose *X* into disjoint subsets 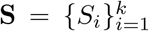 where 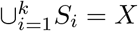.
2. Infer a level-1 network *C*_*i*_ on each subset of taxa *S*_*i*_, giving 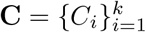.
3. Infer a pairwise dissimilarity matrix *D* on *X*.
4. Utilize InPhyNet to combine the information in *D* and **C** to infer a network 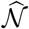 on all taxa in *X*.

Each of these steps is agnostic to the type of input data used or specific method used, so they allow researchers to utilize whatever methods best fit their own use cases. If the networks in **C** and the dissimilarity matrix *D* are inferred in a statistically consistent manner, and the subset decomposition **S** meets certain criteria, we prove in the Statistical Consistency section that such a framework is statistically consistent for inferring the true underlying species network *N* . The main contribution of our work is in developing a novel method for step 4, which we call InPhyNet. One drawback of our approach is that reticulations cannot be inferred between the disjoint subsets **S** constructed in step 1, but this does not mean that interclade or ancestral hybridizations cannot be inferred with InPhyNet. Instead, it means that great care needs to be taken when constructing the subset decomposition. Below, we propose a practical method for performing such a decomposition while maintaining the identifiability of such reticulate relationships.

### Preliminaries

We use the definitions and notation for phylogenetic networks and their components presented in Ané et al. (2024).

#### DEFINITION 1

A *semi-directed phylogenetic network* on taxon set *X* is an unrooted, connected, acyclic, binary, semi-directed graph with nodes *V* = *V*_*L*_*∪V*_*H*_ *∪ V*_*T*_ and edges *E* = *E*_*T*_ *∪ E*_*H*_ with corresponding edge lengths *{l*_*e*_*}* _*e ∈ E*_ and inheritance parameters 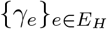. *V*_*L*_ are the network’s set of labeled leaves in bijection with *X, V*_*T*_ are internal *tree vertices* without any incoming directed edges, and *V*_*H*_ are internal *hybrid vertices* with two incoming directed edges and an additional undirected edge. In this paper we assume that all vertices in *V*_*T*_ and *V*_*H*_ have degree 3 whereas those in *V*_*L*_ have degree 1. Additionally, we assume that each *γ*_*e*_ *∈*(0, 1) *{* 0.5*}* and 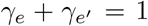where *e* and *e*^*′*^ are directed edges incident to the same hybrid vertex. Tree edges have *γ*_*e*_ = 1.

The restriction *γ*_*e*_ ≠0.5 applies to every network considered in this paper. This restriction guarantees that the major tree and the spanned subnetworks defined below (Definitions 4 and 5) are uniquely defined.

Hereafter, the term *network* refers to a semi-directed phylogenetic network as in Definition 1. Such a network need not contain any hybrid vertices, in which case it is an unrooted binary tree.

#### DEFINITION 2

A *pairwise dissimilarity matrix* is a symmetric matrix with zeros along its diagonal where entry *D*_*AB*_ is a measure of dissimilarity between taxa *A∈X* and *B∈X*. Off-diagonal values are in the interval (0, *∞*) and higher values indicate that *A* and *B* are more dissimilar.

#### DEFINITION 3

A *level-k* network is a semi-directed network in which each biconnected component contains at most *k* hybrid nodes. Counting hybrid nodes in this way relies on the network being binary, as all networks considered in this paper are.

#### DEFINITION 4

The *displayed trees* of network *N* = (*V, E*) are the set of 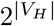 trees obtained by, for each hybrid node *v ∈ V*_*H*_ with incoming hybrid edges *e* and *e*^*′*^, removing either *e* or *e*^*′*^ and fusing edges incident to degree-2 nodes. We denote the set of displayed trees of *N* as disp(*N*). The *major tree* of *N* is the displayed tree obtained by removing *e* if 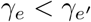, else by removing *e*^*′*^.

#### DEFINITION 5

Let *N* be a network on taxon set *X* and let *Y ⊆X*. Let *T*^*\**^ be the graph obtained from *N* by deleting the minor incoming edge of every hybrid vertex, without fusing edges incident to degree-2 vertices. *T*^*\**^ is a tree that contains every vertex of *N*, so any two taxa are joined by a unique path in *T*^*\**^. Let *U* be the set of vertices lying on each path in *T*^*\**^ between all pairings of taxa in *Y* . The subnetwork of *N spanned by Y* is the graph with vertex set *U* and all edges of *N*, major or minor, whose two endpoints both lie in *U*, after fusing edges incident to degree-2 vertices. A network *N* ^*′*^ on taxa *Y* is said to *agree with* if *Y⊆X* and *N* ^*′*^ is topologically equivalent to the subnetwork of spanned by *Y* .

Examples of Definition 5 on a small network are illustrated in Figure 1. We use Definition 5 in this form because it describes exactly the part of a constraint network that InPhyNet shrinks in step 4 of the algorithm below, and it matches the notion of agreement under which Lemma 1 and Theorem 2 hold.

**Fig. 1.**
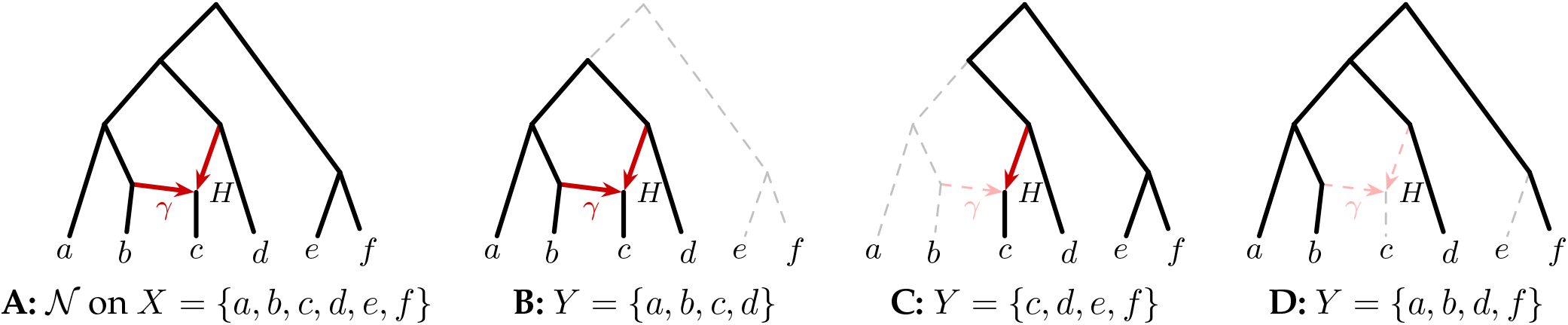
Examples of Definition 5. Networks are drawn rooted for readability. The corresponding semi-directed networks are obtained by suppressing vertices of degree-2. **A:** a network *N* with six taxa *X* = *{a, b, c, d, e, f }* and one hybrid vertex *H*. Tree edges are drawn in black and hybrid edges are drawn in red. **B, C, D:** the subnetworks of *N* spanned by each subset labelled *Y ⊂ X*. Edges included in the spanning networks remain unchanged, while excluded edges are drawn with dashed lines.

#### DEFINITION 6

A *path* from leaf *u* to leaf *w* in network *N* = (*V, E*) is an ordered set of vertices 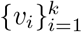 where each *v*_*i*_*∈V* is unique, *v*_1_ = *u, v*_*k*_ = *w*, and an edge exists in *E* that connects either *v*_*j*_ to *v*_*j*+1_ or *v*_*j*+1_ to *v*_*j*_ for *j* = 1, 2, …, *k −*1. We write |*p* |= *k* for the number of vertices in path *p*. Two leaves *u, w* in network are *neighbors* if there exists a path *p* from *u* to *w* in any displayed tree of *N* where |*p*| ⩽ 3, that is, if *u* and *w* form a cherry in some displayed tree. Note in Figure 2 that *u*_min_ and *v*_min_ satisfy this condition in subnetworks (4) and (5) because degree-2 nodes are suppressed in displayed trees, so the path lengths in these cases are 3.

**Fig. 2.**
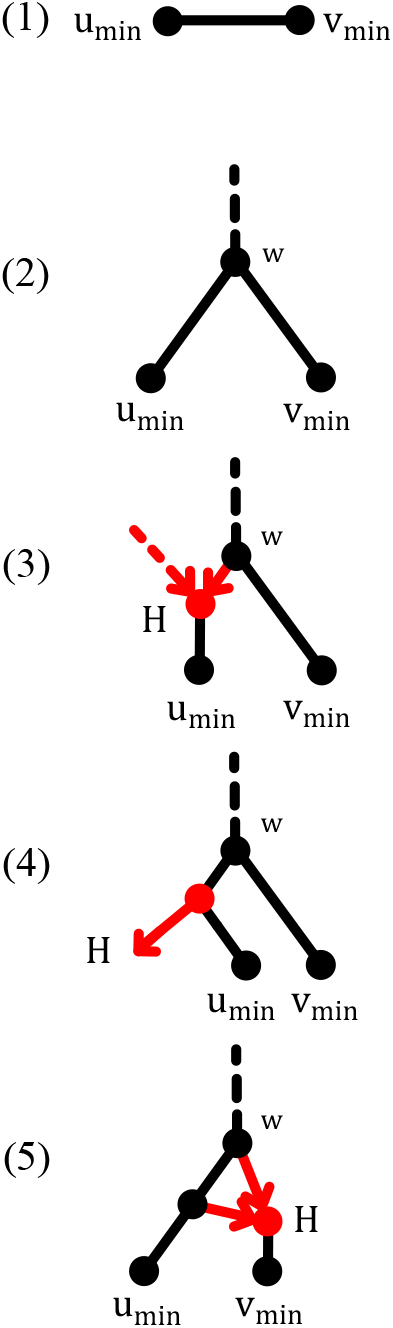
All possible subnetworks in a given constraint network that can be spanned by nodes *u*_min_ and *v*_min_ selected in step (2) of the InPhyNet algorithm. *H* is an arbitrary hybrid vertex in the network and *w* is the node that all non-dashed elements of the subnetwork will be shrunk down to in step (4) of the InPhyNet algorithm. The dashed edge in subnetwork (3) may be present in the constraint network, or may be missing as a result of a subnetwork of type (4) being shrunk previously.

### InPhyNet

The InPhyNet algorithm takes as input a pairwise dissimilarity matrix *D* on *X* along with a set of inferred level-1 networks 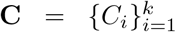 on disjoint subsets of taxa 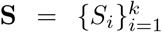 and outputs a network 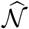 on *X*. We denote this algorithm by 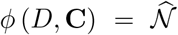. The approach of InPhyNet stems from TreeMerge (Molloy and Warnow, 2019) which itself is derived from Neighbor-Joining (Saitou and Nei, 1987). The algorithm begins with an incomplete network 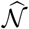 consisting of |*X*| leaves, one for each taxon in *X*, and zero edges (that is, |*X*| isolated vertices). Then, iteratively, the algorithm builds 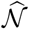 into a full rooted tree via a series of “merge” operations before using the reticulate information contained in the input networks to add reticulations to 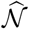.

Finally, the root of the network is suppressed and 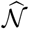 is a semi-directed network. At each iteration, the algorithm can be summarized as follows:

1. Find the set of all pairs of nodes (*u, v*) where either (i) *u, v* are neighbors in every network *C*_*i*_ where both *u, v ∈ C*_*i*_, or (ii) *u, v* do not co-occur in any network *C*_*i*_ *∈* **C**. Call this set *P* . (Algorithm 2)
2. Compute the rate-adjusted matrix *Q* from *D* (as in Saitou and Nei (1987)) to find the most “similar” pair of nodes (*u*_min_, *v*_min_) in *P* .

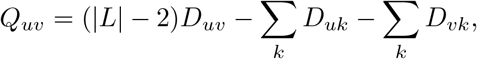

where |*L*| is the current number of nodes under consideration (initially |*X*|, then reduced by 1 after each iteration) and both sums run over all current nodes. Select the pair (*u*_min_, *v*_min_) ∈ *P* with the smallest value of *Q*_*uv*_. In the case of a tie, *u*_min_ and *v*_min_ are chosen randomly. With real-valued dissimilarities estimated from data, exact ties are rare, but when one occurs, different choices can lead to different output networks. (Part of Algorithm 1)
3. Merge *u*_min_ and *v*_min_ in 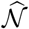via a new shared parent *w*, expanding the inferred network. (Part of Algorithm 1)
4. In each input network *C*_*i*_, shrink the subnetwork spanned by *u*_min_ and *v*_min_ to a single vertex *w*. Record reticulate relationships in this subnetwork so that they may be reflected in 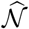 later. (Algorithms 3 and 4)
5. Remove *u*_min_ and *v*_min_ from *D* and add an approximation for their new parent node *w*, with 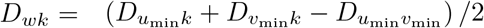 for every other current node *k*. (Part of Algorithm 1)

Informally, InPhyNet builds the network’s backbone by repeatedly joining two nodes before adding reticulations onto that backbone. Pairs of nodes are joined in order of inferred similarity, with the networks **C** providing specific information about the relationships of certain nodes that is assumed to be more trustworthy than the dissimilarity information in *D*. Following this logic, a pair of nodes is only eligible to be joined if every network in **C** containing those nodes validates their siblinghood (they appear as neighbors), or the networks in **C** do not contain information pertaining to the pair of nodes (the nodes never co-occur). Once the set of node pairings has been reduced to *P*, those that are “validated” by the networks **C**, the final decision for that iteration is made according to the inferred dissimilarities in *Q*.

These steps occur |*X*| *−* 1 times, after which the reticulate relationships recorded in step 4 are added to 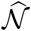 and 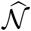 is unrooted, yielding a semi-directed network. The root is removed at this step because its position is not estimated from the data. It is created by the final merge of the algorithm, so its location depends only on the order in which nodes are merged, rather than any biological signal present in the data. For a full worked example with 14 taxa and 3 hybrids, see Appendix 4.

The set of input networks constrains the set of pairs of nodes that are allowed to be merged in step 1, so, hereafter, we refer to these networks as *constraint networks*.

In the case where |**C**| = 2, this process will always successfully conclude so long as the constraints are constructed on disjoint subsets of taxa. However, when |**C**| *>* 2, certain sequences of merges may lead to reduced constraint networks that conflict with one another, leading to |*P*| = 0 in step 1. In this case, the algorithm is adapted as in Molloy and Warnow (2019) as follows:

1. Select *C*_1_, *C*_2_ *∈* **C**.
2. Denote *D*^(12)^ as the portions of *D* that only pertain to leaves in *C*_1_ and *C*_2_.
3. Compute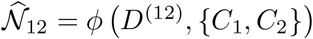.
4. Replace *C*_1_ with 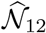 and remove *C*_2_.
5. Repeat until only 1 network remains in **C**. This network is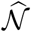.

This ensures that the algorithm terminates with a single network in which every taxon in *X* appears exactly once as a leaf. This procedure is formalized in Algorithm 5. For an example illustrating such a conflict, see Appendix 5. While all of the constraint networks must be level-1 and we present statistical consistency results later for level-1 species networks, this algorithm does not necessarily construct a level-1 network.

### Runtime Complexity

Let |*X*| be the total number of taxa, *r* the total number of reticulations, and *m* the maximum constraint network size. The aforementioned inference framework has three main costs: (i) constraint inference, (ii) pairwise dissimilarity computations, and (iii) the InPhyNet algorithm itself. If the inference method used in (i) has runtime complexity *O*(ℱ (|*X*|)) on |*X*| taxa, then the runtime complexity to infer 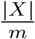straint networks, each of size *m*, is 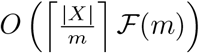. Additionally, (ii) and (iii) have *O*(|*X*| ^2^) and *O*(|*X*| ^3^ + |*X*| *r*) runtime, respectively (see Appendix 3). Thus, the theoretical runtime complexity of such a pipeline is 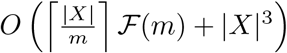. In practice, though, when an accurate inference method is used in (i), the runtime of (ii) and (iii) are negligible. This means that we can expect the runtime of such pipelines to scale *linearly* with respect to the number of taxa in the full network, a claim that we investigate later.

### Recording Reticulate Relationships

For each constraint network *C*_*i*_, if *u*_min_ and *v*_min_ are both contained in the network, the subnetwork that they span can only take the topologies shown in Figure 2. When this subnetwork is shrunk, all vertices shown (except for *w*) and all solid (non-dashed) edges are removed from the constraint network, except for subnetwork (1) which is simply reduced to a single vertex *w*. This leaves the dashed hybrid edge in subnetwork (3) appearing to point to nowhere, but this information is recorded prior to shrinking the subnetwork and is later utilized in Algorithm 3 when reticulations are appended to the network’s inferred backbone.

At the same time, a new vertex *w* is added to the inferred network 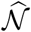 along with edges 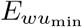 and 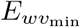 connecting *w* to the other respective vertices. In the cases of subnetworks (1) and (2), no additional work is done. For the remaining subnetworks, information corresponding to the reticulate structures in the subnetwork is logged with respect to the new edges in 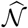 so that this reticulate structure can be reconstructed in 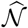. For subnetwork (3) this means that edge 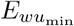is marked as “incoming” for hybrid vertex *H*, for subnetwork (4) 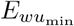 is marked as “outgoing” for *H*, and for subnetwork (5) 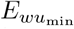 is marked outgoing for *H* and 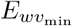 is marked incoming for *H*. In each case, the associated *γ* values in the constraint network are recorded as well.

For each hybrid vertex *H* found in a given constraint network, there will be one edge in 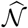 marked as “outgoing” and another marked as “incoming”. In the postprocessing phase of InPhyNet, each of these edges are split in two with a new vertex being added between them, say *Z* and *H*, respectively. The network 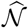 is still rooted at this point, so the edge parent to the hybrid vertex *H* is assigned the respective *γ* value as found in its associated constraint network. Then, a new directed edge is created from *Z* to *H* with an inheritance parameter of 1 *γ*. In cases where a single edge in 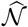 is marked as incoming or outgoing for multiple hybrid nodes, the order in which these markings occurred with respect to the InPhyNet algorithm is stored. Then, the edge is split once for each such respective hybrid with earlier hybrids being assigned lower splits, preserving the ordering observed in the InPhyNet algorithm.

#### Statistical Consistency

##### THEOREM 1.

*Given true species tree T, Neighbor Joining (NJ, Saitou and Nei (1987)) will construct T from a dissimilarity matrix D if, for each pair of true neighbors indexed* (*a, b*) *in D and arbitrary other taxa indexed* (*x, y*) *in D*, 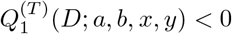 *and* 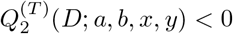.

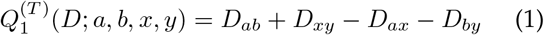

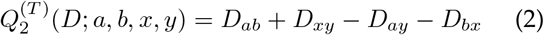

*Proof*. Presented in Appendix C of Saitou and Nei (1987). □

Note that Theorem 1 makes no assumption on the underlying data used to generate *D*, but instead asserts a property that *D* must uphold in order to construct *T* . We examine whether such a matrix *D* constructed under the average gene tree internode distance (AGID) metric upholds this property under the MSNC model later.

##### DEFINITION 7

Let *G*_*n*_ be input data of size *n* (for example, *n* gene trees or an MSA with *n* base pairs) generated under the true species network *N* on taxon set *X*, and for *S ⊆X* let *G*_*n*_|_*S*_ denote the data restricted to the taxa in *S*.

a. A method *M*_*D*_ for computing a pairwise dissimilarity matrix *D* = *M*_*D*_(*G* _*n*_) on *X* is *statistically consistent* for *N* if there exists a displayed tree *T* ∈ disp(*N*) such that, for each pair of true neighbors (*a, b*) in *T* and arbitrary other taxa (*x, y*):

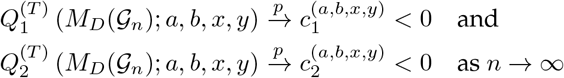

where 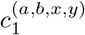and 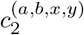 are arbitrary constants that may differ depending on the choices of *a, b, x, y*.
b. A method *M*_*C*_ for inferring level-1 networks is *statistically consistent* for *N* if:

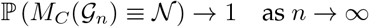

where *≡* indicates topological isomorphism between two networks.
c. An estimator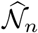 computed from *G*_*n*_ is *statistically consistent* for *N* if 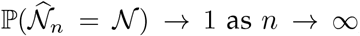 as *n→ ∞*, where equality means equality of semi-directed topologies.

We now establish the conditions under which the network constructed by InPhyNet and the true network share a common displayed tree. InPhyNet works by first constructing a tree and then appending hybrid edges onto this tree, so such a condition is necessary to later establishing when InPhyNet constructs the exact true *network*.

##### Lemma 1

Given the true species network *N*, dissimilarity matrix *D*, and constraint networks **C**, if *D* satisfies Equations 1 and 2 for some *T* ∈ disp *(N* )(the set of displayed trees of *N*) and all constraint networks agree with *N*, then *T* ∈ disp (*ϕ*(*D*, **C**)) where *ϕ*(*D*, **C**) is the network inferred by InPhyNet.

*Proof*. Notice that the final inferred network of InPhyNet is constructed by adding reticulate edges to a tree phylogeny, call that phylogeny 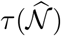. For each constraint network *C*_*i*_ *∈* **C**, because each *C*_*i*_ agrees with *N*, any reticulations that are suppressed in *N* to obtain *T* can similarly be suppressed in *C*_*i*_, if they exist in *C*_*i*_, to obtain a displayed tree of *C*_*i*_ that agrees with *T* . It follows that at each iteration of InPhyNet, the pair ofnodes *u*^*\**^, *v*^*\**^ for which the rate-adjusted matrix *Q* is minimized will belong to *P* because, by Definition 6, they will be considered neighbors in every *C*_*i*_ where they both appear. Thus, at each iteration, the global pair of nodes minimizing *Q* will be merged, which is exactly the NJ algorithm, so 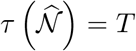. Whether 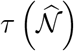 is the major tree of 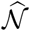 or simply some minor tree depends on the specific *γ* values inferred in the constraint networks C. □

Here, it should be noted that a network is not encoded by its set of displayed trees (Zhu and Degnan, 2017). This fact poses significant methodological challenges when it comes to inferring networks under the MSNC. In a pipeline utilizing InPhyNet, these challenges fall on the methods used to compute InPhyNet’s inputs rather than InPhyNet itself.

Next, we utilize Lemma 1 to establish the conditions under which InPhyNet constructs the exact true network.

##### Theorem 2.

*Given the true species network N, constraint networks* **C** *on non-overlapping sets of taxa* **S**, *and dissimilarity matrix D, the network inferred by InPhyNet* 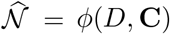 *is exactly N if the following are true:*

1. *D satisfies Equations 1 and 2 for some T ∈ disp*(*N* ),
2. *all C*_*i*_ *are level-1 and agree with N (see Definition 5), and*
3. *for each hybrid vertex in N, at least one subset in* **S** *contains at least one individual from each of the clades A, B, C, D, and H as shown in Figure 4*.

*Proof*. From assumption (1) and (2), if InPhyNet’s initial merging proposals are always accepted, then according to Lemma 1 *T ∈* disp 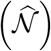. Then, all that remains to be shown is that the reticulations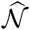 in are placed in the same positions as they appear in *N*.

For each hybrid vertex *v ∈ V*_*H*_ with incoming hybrid edges *e* and *e*^*′*^, we refer to the hybrid vertex’s *reticulation* as whichever edge *e* or *e*^*′*^ is *not* displayed in *T* . Then, for each reticulation *r*_*j*_, the various components of the cycle formed by *r*_*j*_ in *N* and their corresponding components in the constraint network that contains *r*_*j*_ are identified in Figure 4. Notice that by assumptions (2) and (3) the cyclic structure shown in Figure 4 will also appear in at least one constraint network. For the remainder of the proof, we refer to the corresponding vertices in this cyclic structure as they appear in the constraint networks as *Ã*, 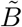, 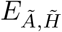, and so on.

Notice that in order for a hybrid node to be created in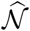, two nodes in some constraint network *C*_*k*_ must be neighbors connected by a path that either (a) contains a hybrid node, or (b) contains a node with a directed edge pointing towards a hybrid node. We have shown 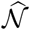 that will display *T*, so it follows that for each 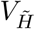 one of its incoming directed edges 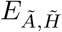 or 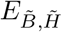 will be traversed in this path and therein be placed correctly in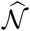. Without loss of generality, assume that this edge is 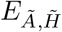 .

Then, notice that nodes *V*_*B*_ and *V*_*C*_ would not both appear in a displayed tree of *N* because one of them will be suppressed in the displayed tree along with *V*_*H*_ . If clades *B* and *C* are merged after each of their members are merged, though, such a node can be created in the inferred network by splitting edge *E*_*B*_ (as in Figure3). When 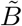 and 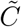are merged, 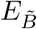 is traversed, where-after it is created in 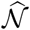nd marked as outgoing to *H*. Similarly, when *Ã* and 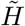are merged, 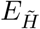is traversed, whereafter it is created in 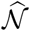 and marked as incoming to *H*. Then, in the post-processing step of InPhyNet, the exact aforementioned process needed to place the hybrid properly of splitting these edges is conducted.

Thus, it follows that *V*_*B*_ and thereby *E*_*B,H*_ will be placed correctly in 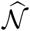 if (i) all members of 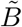 and 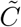 merge with one another before they do with any other taxa in the constraint network, and (ii) when 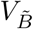and 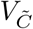are merged, edges 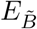 and 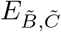 occur in the subnetwork spanned by 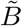and 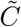. Condition (i) follows from assumption (1). Given that we took *Ã* and 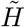to have been merged first without loss of generality, any merging of 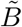and 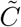must go through edges 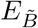and 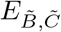, so condition (ii) is satisfied as long as 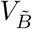 and 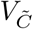 are neighbors in the constraint network. The presence of these vertices in the constraint network is given by assumption (3), and their status as neighbors is given by assumption (2). It follows, then, that each hybrid vertex and its respective hybrid edges will be placed correctly in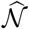 . □

**Fig. 3.**
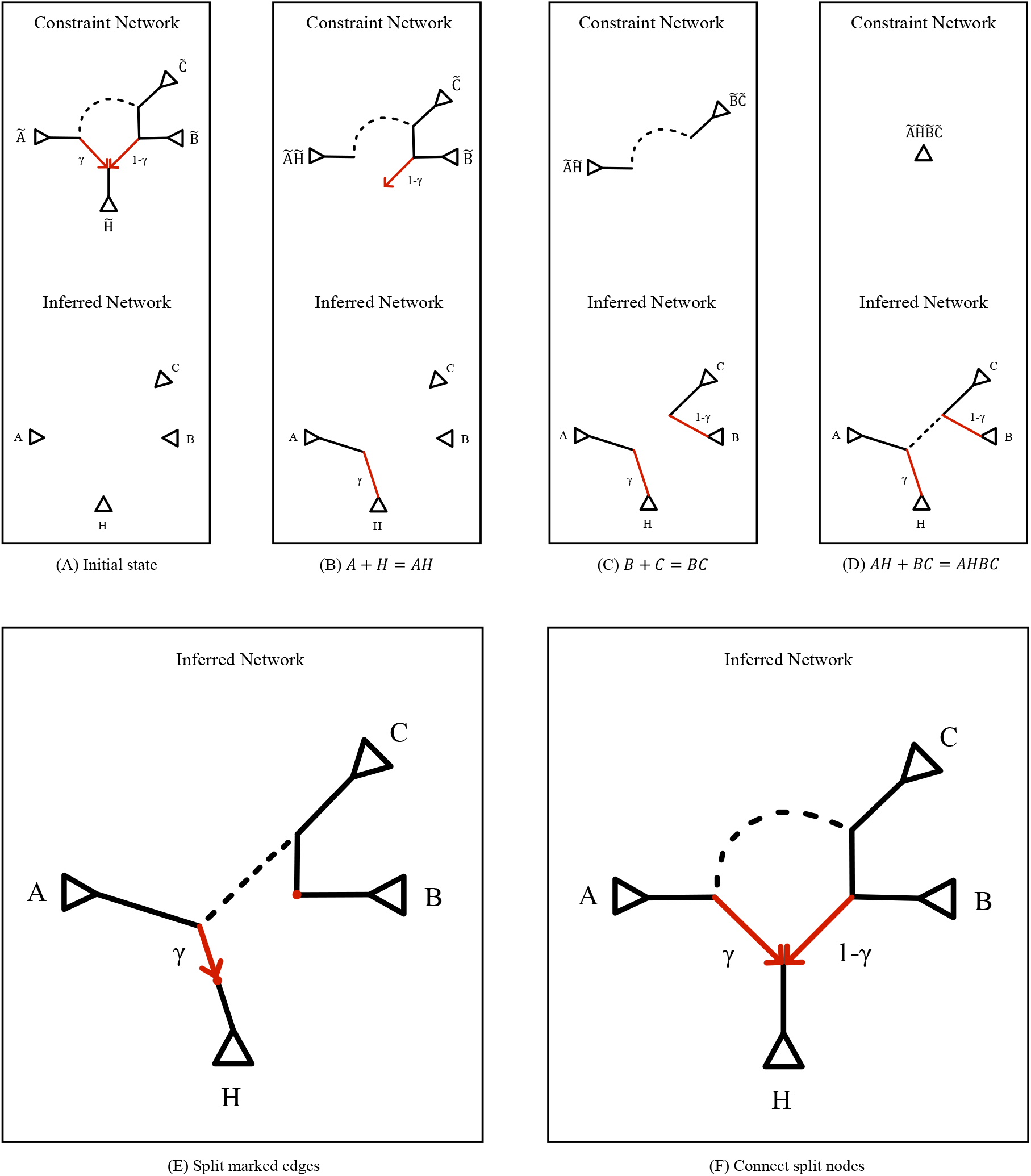
**A:** the initial relevant state of the algorithm at which point all taxa within each clade have been merged. **B:** nodes *A* and *H* are merged. The path *p*^*\**^ connecting them in the constraint network traverses one reticulation, so its corresponding edge is marked in the inferred network. **C:** nodes *B* and *C* are merged where *p*^*\**^ traverses a node with an outgoing reticulation, so this reticulation is also recorded in the inferred network. **D:** there may be zero, one, or multiple mergings along the dashed edge before this step occurs, but those do not affect how *A, B*, and *C* are ordered around *H*. **E:** the edges that correspond to reticulations are split. The edges are split differently based on whether they correspond to an incoming or outgoing reticulation. **F:** a reticulation is drawn from the red node above *B* to the hybrid node above *H* to complete the inferred network.

**Fig. 4.**
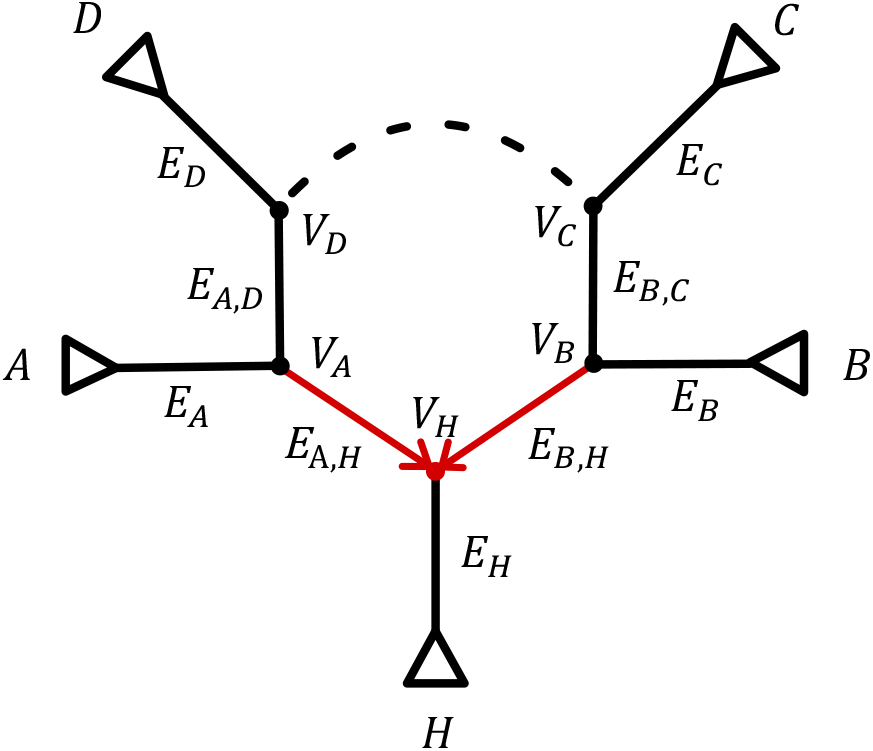
Cycle corresponding to hybrid node *V*_*H*_ in the true species network with labelings corresponding to those used in Theorem 2. This structure is also displayed in any constraint that meets condition 3 of Theorem 2 but with labels corresponding to the relevant subsets of each clade *Ã ⊆ A*, 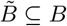, and so on.

Theorem 2 comes with strong assumptions, but this is due to the strength and guarantee of its outcome. In practice, these assumptions may not be strictly required for perfect or near-perfect method performance. For example, assumption (1) requires that *D* takes a very strict form. For the purposes of Theorem 2 this ensures that siblinghood relationships represented in the inferred network will be correct even when a given pair of taxa do not appear together in any of the constraints. However, in practice, there is significant room for error in *D* so long as the constraint networks are accurately inferred, because many incorrect siblinghood relationships implied by *D* may be disallowed by the relationships in the constraint networks.

Additionally, assumption (2) is required for strict isomorphism between the true and inferred networks, but small topological deviations between the true and inferred constraint networks will naturally only translate to small deviations between the true and inferred species networks. When the focus of analysis is on 100s of taxa, these granular differences will typically be contained within a given clade and will thus have a negligible impact overall.

Finally, assumption (3) is perhaps the strongest as it cannot be known in practice. The necessity of this assumption is highly informative, though, as it provides a clear goal and direction for the subset decomposition phase.

Now, we apply Theorem 2 in the context of a statistically consistent framework to show that a statistically consistent pipeline utilizing InPhyNet is theoretically possible.

##### Corollary 1

Call *M*_*C*_ 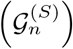 a network inferred on the set of taxa *S* using only the portion of data in *G*_*n*_ that pertains to *S*. Given input data *G*_*n*_ of size *n* on taxa *X* generated under true species network *N* with disjoint subset decomposition 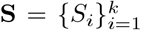, if *M*_*C*_ and *M*_*D*_ are both statistically consistent in the sense of Definition 7 and **S** satisfies condition (3) in Theorem 2, then the esti-mator 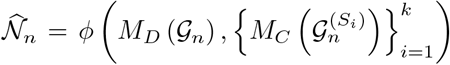is statis-tically consistent with respect to *n* for the true species network *N* . In other words,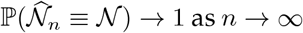

*Proof*. Write 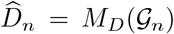 for the estimated dissim-ilarity matrix and 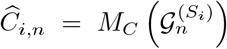, so that 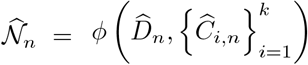 . Condition (3) of Theorem 2 holds by assumption, so by Theorem 2, 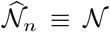 whenever 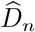 satisfies condition (1) and every 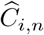 agrees with *N* . We bound the probability of each of these events separately (Steps 1 and 2) and then combine the bounds (Steps 3 and 4).

*Step 1 (Dissimilarity Matrix)*

Let *T∈* disp(*N* ) be the displayed tree given by Definition 7(a). Index the sets of four taxa in *X* by 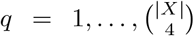 and let *A*_*q,n*_ be the event that 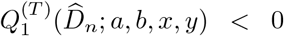 and 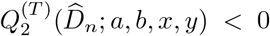 for each choice of true neighbors (*a, b*) in *T* and other taxa (*x, y*) within the *q*-th set. By Definition 7(a), each of these finitely many quantities converges in probability to a negative constant, so p(*A*_*q,n*_) *→* 1. That is, for every *ϵ*_*q*_ *>* 0, there is an *n*_*q*_ such that p (*A*_*q,n*_) ⩾ 1 *− ϵ*_*q*_ for all *n* ⩾ *n*_*q*_. If every event *A*_*q,n*_ occurs, then *D*_*n*_ satisfies Equations 1 and 2 for *T*, which is condition (1). The probability that at least one *A*_*q,n*_ fails is at most the sum of the probabilities that each of them fails, so the probability that *D*_*n*_ satisfies condition (1) is given by:

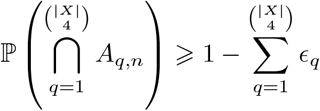

*Step 2 (Constraint Networks)*. Applying Definition 7(b) to the data 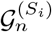 on the taxa *S*_*i*_, for every 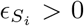 there is an 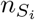 such that, for all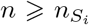, the probability that 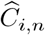 agrees with *N* is at least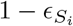.

*Step 3 (Combining Steps 1 and 2)*. Let 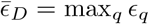and 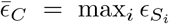. Assuming that 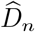 and 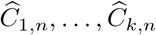 are independent, the probability that all of the events in Steps 1 and 2 occur is the product of their probabilities. Hence, for all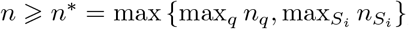,

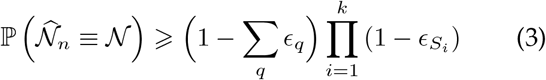

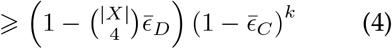

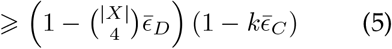

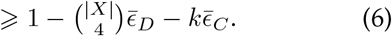

Equation 3 uses steps 1 and 2. Equation 5 is Bernoulli’s inequality ((1 *−t*)^*k*^ ⩾ 1 *−kt* for 0 ⩽ *t* ⩽ 1). Equation 6 expands the product and drops the nonnegative term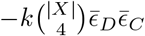.

*Step 4 (Conclusion)*. Given any *ϵ >* 0, choose 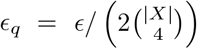 for every *q* and 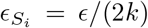 for every *S*_*i*_. Then the right-hand side of Equation 6 equals 1 *− ϵ*, so 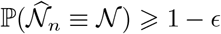 for all *n* ⩾ *n*^*\**^. Since *ϵ* was arbitrary,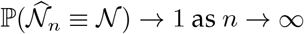. □

## Performance Study

### Overview

We quantify the estimation error of the inference framework presented above with various amounts of input data (number of gene trees or sequence length), numbers of taxa, levels of incomplete lineage sorting (ILS), and sizes of constraint networks (*m*). See Table 1 for all parameter values. We evaluated the topological accuracy of the framework by calculating the unrooted hardwired cluster dissimilarity (HWCD, Huson et al., 2010) between the inferred species network and the true species network, namely HWCD(*N*,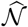 ). These calculations were done using the `hardwiredclusterdistance` function in the PhyloNetworks Julia package (Solís-Lemus et al., 2017). HWCD is a dissimilarity measure defined for *rooted* networks, but can be calculated on unrooted networks by finding the rootings of each network that minimize the rooted HWCD. Additionally, we evaluated the runtime of the framework across each simulation setting. The original data used in these simulations can be found on Dryad, and the scripts used to analyze this data can be found at github.com/NathanKolbow/InPhyNetSimulations.

**Table 1.** Simulation parameters and values used in the simulation study. Low and high ILS correspond to average network branch lengths of 1.5 and 0.15, respectively. Ten replicates were conducted for each parameter combination.

| Parameter | Values |
| --- | --- |
| Number of taxa | 25, 50, 100, 200 |
| Number of base pairs | 100, 1000 |
| Number of loci | 100, 1000 |
| Maximum constraint size ($m$) | 10, 20 |
| Incomplete lineage sorting (ILS) | low, high |

### Simulated Data

Specific settings and parameters used for each of the software mentioned below can be found in Appendix 2. The ground truth species networks were simulated according to a birth-death-hybridization model with the SiPhyNetwork package in R (Justison et al., 2023). This package simulates networks of arbitrary level, so in each case networks were simulated with fixed parameters until a level-1 network was obtained. Then, gene trees were simulated according to the multispecies network coalescent model using the PhyloCoalSimulations package in Julia (Fogg et al., 2023). Finally, multiple sequence alignments (MSAs) were simulated with seq-gen (Rambaut and Grass, 1997) under the F81 model of evolution (Felsenstein, 1981) with state frequencies *π*_*A*_ = *π*_*T*_ = 0.3 and *π*_*C*_ = *π*_*G*_ = 0.2.

### Framework Methods

We used SNaQ (Solís-Lemus and Ané, 2016), PhyloNet’s maximum likelihood option (PhyloNet-ML) (Yu et al., 2014), PhyloNet’s maximum pseudolikelihood option (PhyloNet-MPL) (Yu and Nakhleh, 2015), and Squirrel (Holtgrefe et al., 2025) to infer the constraint networks. It should be noted, though, that Squirrel is not a coalescent-aware method, and so these simulations are explicit model violations. Analyses with PhyloNet-ML were restricted to only using *m* = 10 due to prohibitively long runtimes. In addition, eight analyses utilizing PhyloNet-ML with *m* = 10 were excluded due to a failure to complete within 1 month.

The average gene tree internode distance (AGID, see Definition 8 for a formal definition) metric was used to compute the pairwise dissimilarity matrix *D* in each simulation, and the methodology presented in Materials and Methods was used for subset decomposition. We established in the section Statistical Consistency that this framework can be statistically consistent when *M*_*C*_ and *M*_*D*_ are statistically consistent and subset decomposition is done in an appropriate manner, but we do not claim that this specific implementation of the framework meets these criteria. In particular, AGID is *not* statistically consistent in general in the sense of Definition 7 (see Appendix 1).

#### Subset Decomposition

The primary goal of a subset decomposition method in this framework is to (a) identify areas where hybrids may be present, and (b) choose subsets such that the conditions in Theorem 2 are met. In doing so, we infer the *tree of blobs* of the species network (Allman et al., 2023). The tree of blobs of a network is a treelike structure where each of the network’s maximally biconnected components, called *blobs*, are shrunk to individual vertices. This structure is useful for subset decomposition because it provides information about the locations of cycles in a network and can be inferred much more efficiently than a network. We implement subset decomposition as follows:

1. Infer the tree of blobs *T* of the true species network with TINNiK (Allman et al., 2024).
2. For each blob *B*_*i*_ in *T*, the blob is connected to edges *E*_1_, *E*_2_, …, *E*_*k*_. For each of these edges, imagine splitting *T* in half at that edge, and randomly selecting a single leaf node that is *not* in the same split half as the blob center. Conduct this process for each edge *E*_1_, *E*_2_, …, *E*_*k*_, and add these *k* leaves to a set *L*_*i*_ associated with blob *B*_*i*_.
3. Infer a species tree 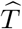 on the full set of taxa with ASTRAL (Zhang et al., 2018).
4. For each set *L*_*i*_, remove all taxa in *L*_*i*_ from 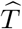 except for the first entry of *L*_*i*_, denote this taxon as 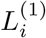. This taxon will be used to represent the set *L*_*i*_.
5. Select the edge in 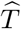 that splits 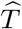 into equally sized subtrees (or as close to equally sized as possible). Repeat this recursively on the resulting subtrees until each subtree contains no more than *m* leaves.
6. Find each representative taxon 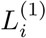 and its corresponding subtree and expand 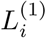 into all of *L*_*i*_.

The taxa contained in each subtree now correspond to the subset decomposition *S*_*i*_.

We found that when a tree of blobs is inferred with more than 50 taxa, step (1) is computationally infeasible. In this case, we identify blobs in batches by first inferring a species tree with ASTRAL and then decomposing the set of taxa into subsets of size 50 in both manners depicted in Figure 5. Then, steps (1) and (2) are conducted on every such subset of size 50 constructed in this manner, generating the collection of sets *L*_*i*_ which is then used for each of steps (4)–(6). This approach allows us to identify hybridization events in the true species network both when they occur between distant and local lineages.

**Fig. 5.**
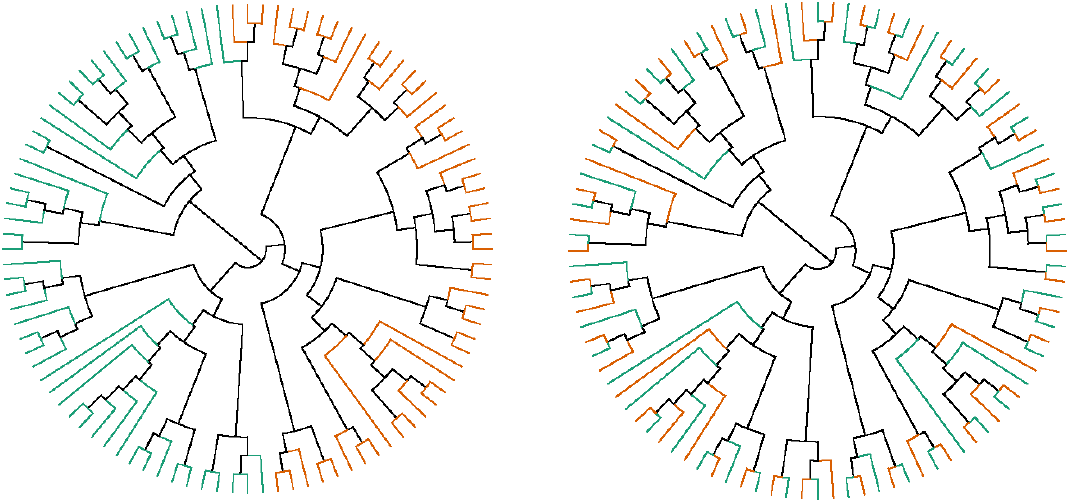
An illustration of a species tree 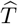 inferred on 100 taxa being split into subsets of 50 individuals (one subset in green, the other in orange) in two different ways, as described in Subset Decomposition. **Left:** two subsets of 50 taxa split such that the majority of any given clade tends to appear in the same subset. **Right:** two subsets of 50 taxa split such that distant taxa tend to appear in the same subset.

#### Inference with SNaQ

Analyses with the SNaQ method were conducted with the SNaQ package in Julia (Kolbow et al., 2026). SNaQ takes estimated quartet concordance factors as input, which require estimated gene trees to be computed. So, we inferred gene trees using IQ-TREE (Minh et al., 2020) and used ModelFinder for model selection. Further, each instance of SNaQ requires a maximum number of hybrids (`hmax`) to be specified before a network can be inferred. In practice, for a fixed set of taxa, a set of networks with ascending `hmax` values are inferred before model selection is conducted. This significantly increases the computational burden of each simulation, so instead, for a given subset of taxa *S*_*i*_ from true species network *N*, we selected `hmax` to be the number of hybrids present in the subnetwork of *N* spanned by *S*_*i*_.

#### Inference with Squirrel

Analyses with the Squirrel method were conducted with the `physquirrel` package in Python. Dense sets of quarnets were inferred from the respective MSA with the `delta heuristic` method, and we inferred level-1 networks from these dense sets of quarnets with the `squirrel` method.

### Results

#### Accuracy

Output error, measured as the HWCD between the inferred species network and the true species network, tended to decrease as the number of gene trees and number of base pairs increased in the simulations (Figure 7). We observed comparable accuracy when utilizing SNaQ, PhyloNet-ML, and PhyloNet-MPL in tandem with InPhyNet, but observed significantly worse performance when utilizing Squirrel. In cases with low ILS, median error across all simulation settings was near zero with all methods except for Squirrel. Simulations with high ILS did not perform as well, however. We found that the maximum constraint network size (*m*) did not have a noticeable impact on accuracy in any simulations with any underlying inference method.

We observed a very strong dependence between the total error in the input networks given to InPhyNet and the output error of the network output by InPhyNet (Figure 6). This indicates that InPhyNet is highly robust to error in the dissimilarity matrix, but highly sensitive to error in the input networks.

**Fig. 6.**
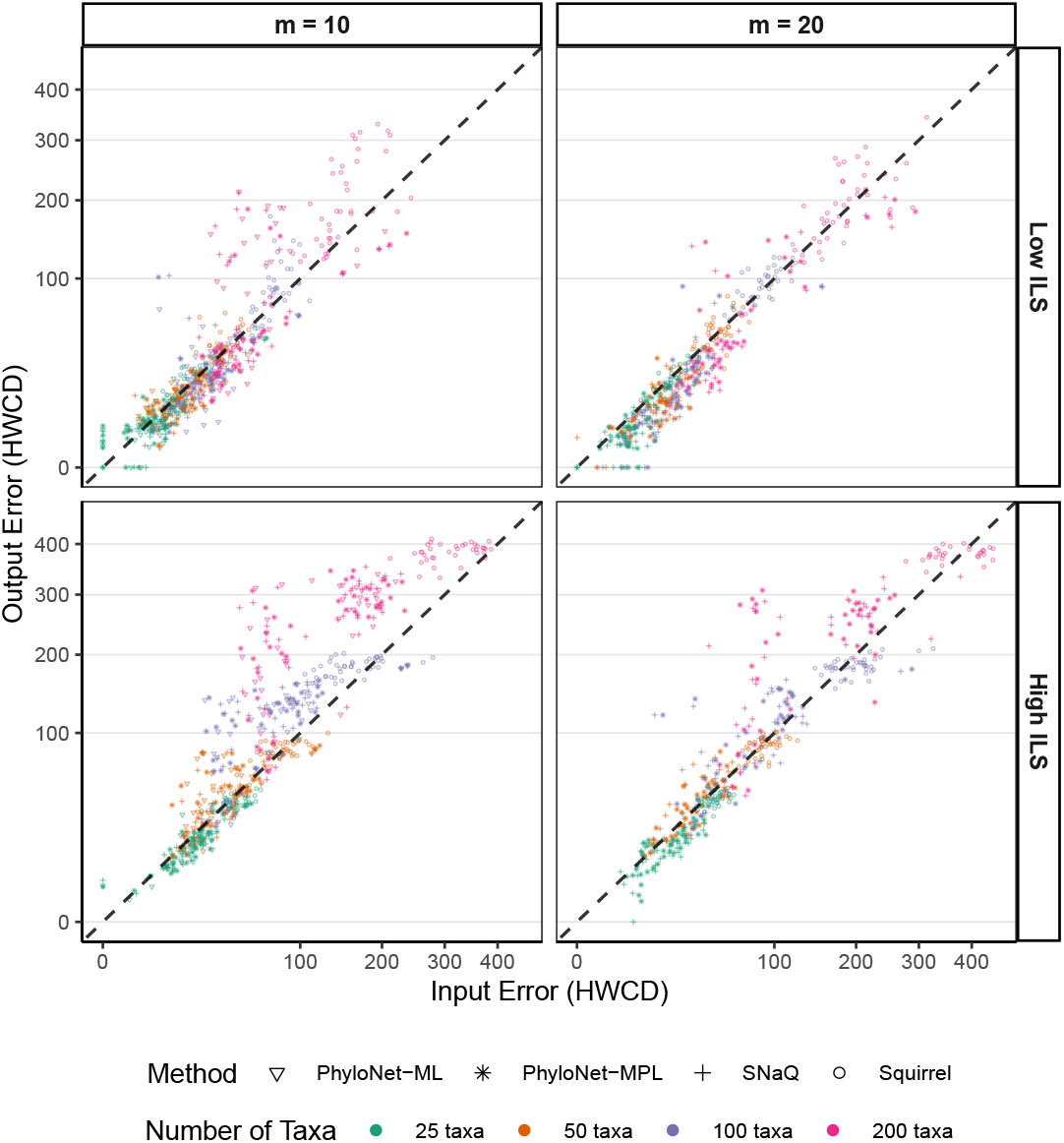
Output error of the species network inferred by InPhyNet in unrooted hardwired cluster dissimilarity (HWCD) vs. the total input error to InPhyNet. Input error is calculated by summing the inference errors of each constraint network used as input to InPhyNet. The inference error of each constraint is calculated by taking the HWCD between the constraint network and the subnetwork of the true species network that is spanned by the taxa contained in the constraint network.

**Fig. 7.**
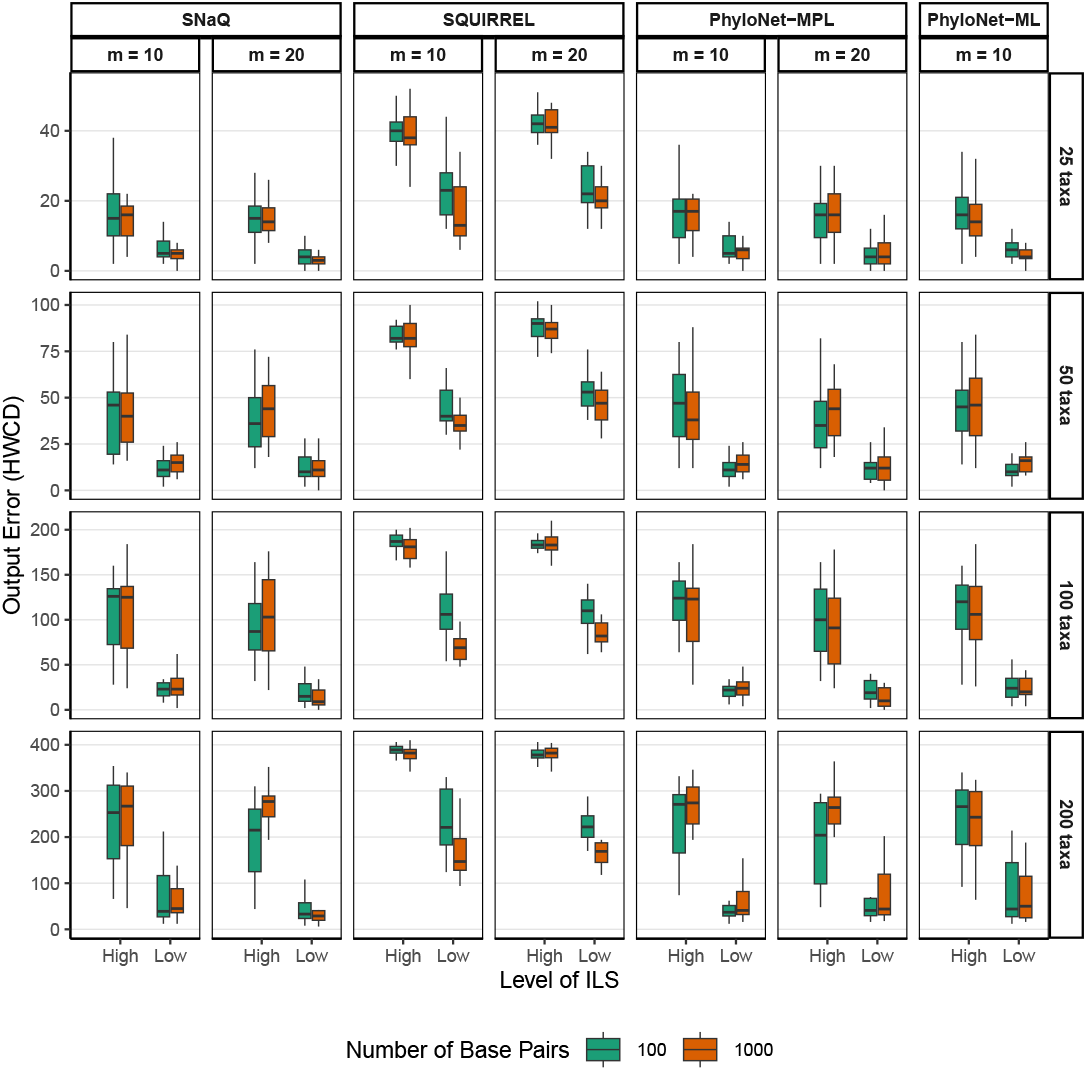
Output error of the pipeline across various simulation settings in hardwired cluster dissimilarity (HWCD). *m* denotes the maximum number of tips included in each constraint network. The labels running along the rightmost subplots indicate the number of tips in the true species network.

#### Runtime

For a fixed maximum constraint network size *m*, total runtime increased approximately linearly with respect to the number of taxa in the full network, |*X*| (Figure 8). Runtimes were significantly longer for simulations with *m* = 20 than their counterparts with *m* = 10. Specifically, simulations utilizing Squirrel, SNaQ, and PhyloNet-MPL with *m* = 20 took 2.6, 10.9, and 8.9 times longer than their counterparts with *m* = 10. Inference was only conducted with PhyloNet-ML with *m* = 10. All simulations utilizing Squirrel took at most 10 minutes to complete, whereas simulations with SNaQ and PhyloNet took on the order of tens or hundreds of hours to complete, depending on *m*.

**Fig. 8.**
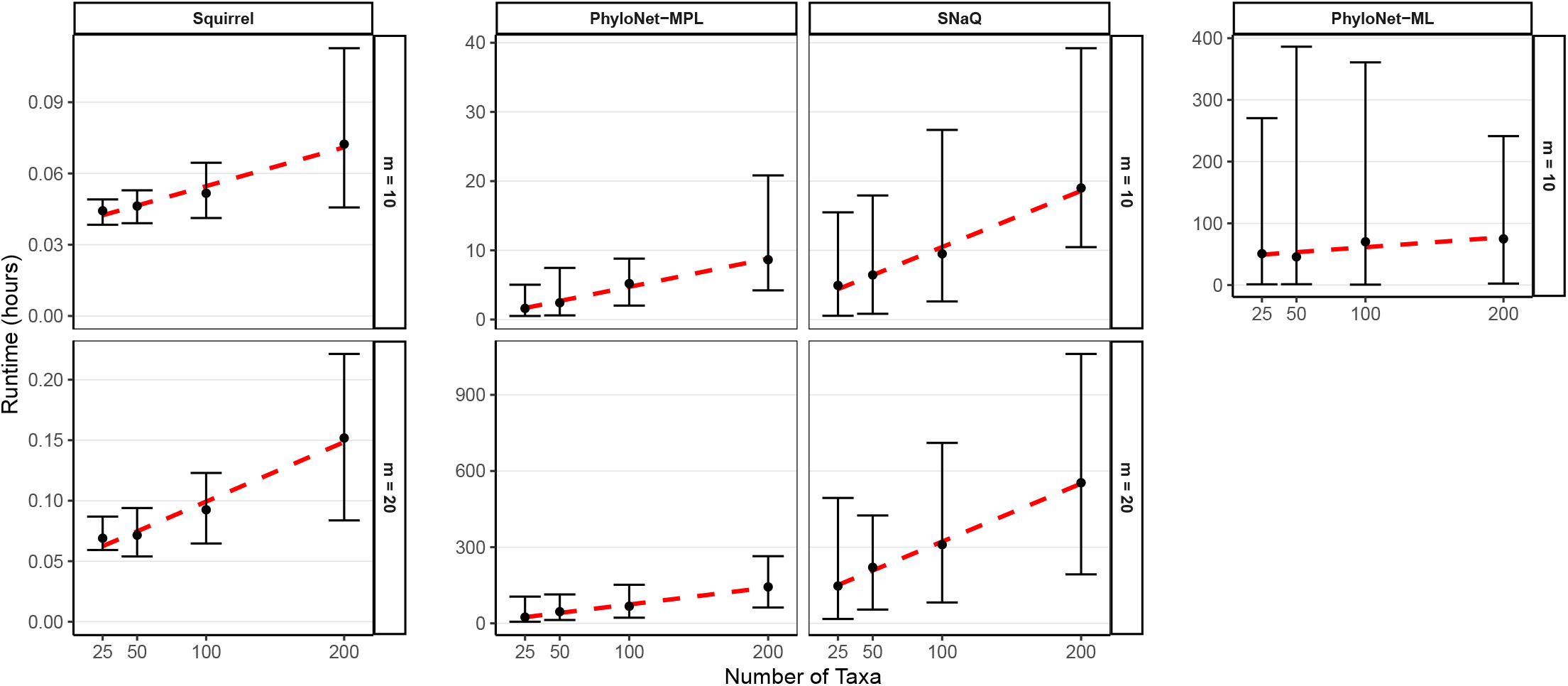
95% confidence intervals for total runtime (in hours) to infer constraint networks, estimate the pairwise dissimilarity matrix *D*, and infer the final network with InPhyNet across various inference methods in settings with low levels of incomplete lineage sorting (ILS). Each column of subplots corresponds to a different inference method whereas each row corresponds to different maximum constraint network sizes (*m*).

## Phylogeny of Green Plants

We re-analyze the phylogeny of 1,158 green plants presented in One Thousand Plant Transcriptomes Initiative (2019). Phylogenetic tree analyses led to observed quartet concordance frequencies that conflicted with what would be expected under the inferred model. Specifically, there was significant conflict between concordance factors involving four major gymnosperm clades and between concordance factors involving four major fern clades. These areas of the phylogeny have previously been shown to exhibit evidence of reticulate evolution, so we utilized InPhyNet to incorporate such reticulations into the phylogeny of green plants. Portions of the tree phylogeny outside of the gymnosperms and ferns were well resolved, so we utilized the subnetwork of the phylogeny inferred in One Thousand Plant Transcriptomes Initiative (2019) spanned by all non-gymnosperm and non-fern species as a single constraint network in InPhyNet. Constraint networks for the remaining species were inferred using the aforementioned inference pipeline with a maximum constraint size of *m* = 20. The subset decomposition step was done manually to ensure that all apparent instances of reticulate evolution would be identifiable by SNaQ. For further details, see Appendix 6.

## Results

The network of green plants that we inferred is presented in Figure 9. While this species network generally agrees with previously inferred phylogenies (One Thousand Plant Transcriptomes Initiative, 2019; Bowe et al., 2000; Liu et al., 2022; Wickett et al., 2014), it also provides new insights into regions of persistent discordance. For example, the placement of asterids, rosids, Caryophyllales, and Saxifragales remains unresolved, which is a particularly consequential issue because together these clades represent approximately 75% of all angiosperm species (Wang et al., 2009). Other recent large-scale phylogenomic studies (Zuntini et al., 2024; Kates et al., 2024) have also highlighted ongoing uncertainty in these relationships, underscoring that this lack of resolution is not unique to our re-analysis. In addition, the network recovered reticulations that help reconcile competing hypotheses regarding the placement of Order Gnetales within gymnosperms, and highlights evidence of reticulate evolution within Pinaceae and Polypodiidae. In the following subsections, we examine each of these cases in detail, drawing on prior research to substantiate the inferred reticulations.

**Fig. 9.**
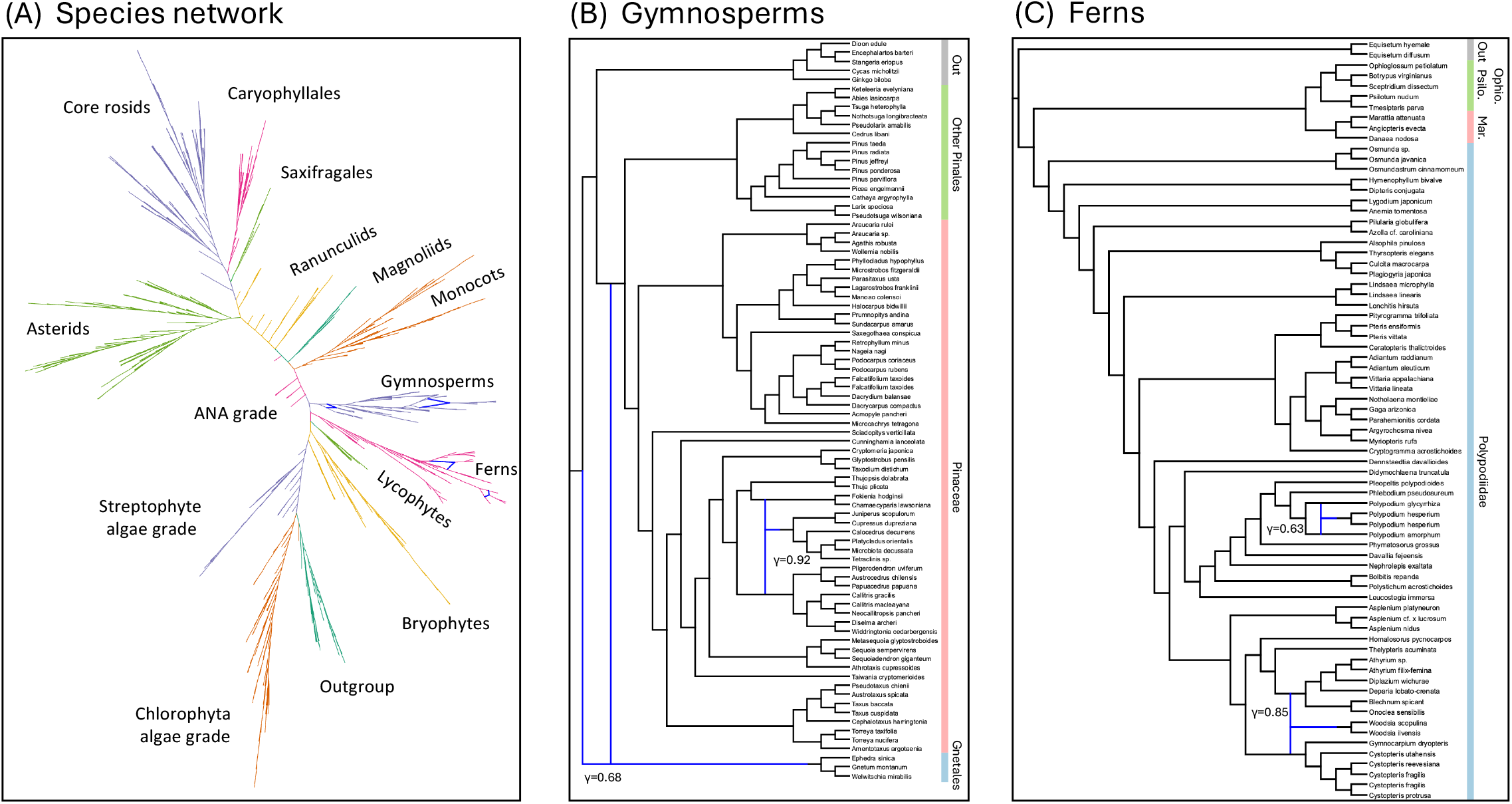
**A:** Semi-directed phylogenetic network of green plants obtained using InPhyNet. Various colored branches signify different clades within the phylogeny. Branch lengths are not scaled. **B:** Magnified portion of the phylogeny corresponding to gymnosperms. The reticulation ancestral to Gnetales provides support to two controversial placements of Gnetales within the phylogeny of gymnosperms. The reticulation inferred within Pinaceae is corroborated by previous research into the histories of each of its descendant species (Farhat et al., 2019, 2022; Zonneveld, 2012). The inheritance probability (*γ*) for each reticulation is noted in the network. **C:** Portion of the phylogeny corresponding to ferns. Previously reported phylogenetic trees for this group revealed significant discordance between the four major clades shown here. However, SNaQ and InPhyNet inferred no reticulations between these clades. Our analyses resulted in two reticulations within Polypodiidae that are congruent with previous studies (Sigel et al., 2014; Smith et al., 2006; Shao et al., 2015).

### Gnetales

The placement of Gnetales in the phylogeny of gymnosperms is controversial. Some research suggests that they are sister to Pinaceae (the “Gnepine” hypothesis), others suggest that they are sister to conifers (the “Gnetifer” hypothesis) and others still suggest that they are sister to cupressophytes (the “Gne-cup” hypothesis) or to all conifers together (Bowe et al., 2000; Soltis et al., 2002; Ran et al., 2018; Chaw et al., 2000; Zhong et al., 2011; Braukmann et al., 2009; Magallón and Sanderson, 2002; Coiro et al., 2022). Treelike analyses of these species have led to significant conflict between these hypotheses, but our analyses reconcile them by providing partial support for both the Gnetifer and Gnepine hypotheses. However, because the inferred phylogeny of Gnetales is known to be highly sensitive to the type of data analyzed and the handling of systematic biases (Zhong et al., 2011), this result should be regarded with caution and warrants further investigation.

### Pinaceae

High rates of polyploidy have been identified in genus *Juniperus* (Farhat et al., 2019), and similar evidence has been found in *Cupressales* and *Platycladus* (Farhat et al., 2022). We are unaware of such direct evidence for *Calocedrus decurrens, Microbiota decussata*, or *Tetraclinis spp*., but research suggests that these three species are closely related to *Platycladus orientalis* (Zonneveld, 2012), corroborating their placement in this phylogeny.

While SNaQ does not directly model polyploidization, PhyloNet (Wen et al., 2018), a similar likelihoodbased approach, has been shown to accurately infer polyploidy events in genus *Fragaria* despite not explicitly modeling polyploidization (Yan et al., 2022). Accordingly, these reticulations in the inferred network may reflect hybridization events, including allopolyploidy, but they could also arise from other sources of conflict, and their interpretation remains uncertain.

*Genera Polypodium and Woodsia* The reticulation inferred in *Polypodium* is well-founded as *Polypodium hesperium* is an allotetraploid and its progenitors are precisely *Polypodium amorphum* and *Polypodium glycyrrhiza* (Sigel et al., 2014). The reticulation in *Woodsia* can also explain previous works that have inferred either a multifurcation between genera *Woodsia, Athyrium, Diplazium, Blechnum, Onoclea*, and other ferns (Smith et al., 2006), or that have otherwise found generally weak support for treelike backbones between these species (Shao et al., 2015).

## Discussion

### Performance Study

Our performance study demonstrates that the accuracy of InPhyNet is strongly associated with the accuracy of the input networks (see Figure 6). The magnitude of this relationship, however, depends markedly on the number of taxa in the network and on the value of *m*. In particular, simulations with fewer taxa more frequently produced output errors that were less than or equal to the corresponding input errors. For datasets of 25, 50, 100, and 200 taxa, this outcome was observed in 82.3%, 61.4%, 52.4%, and 39.2% of simulations, respectively. This trend also seems to depend on the chosen value of *m* and the level of ILS present in the true network. This outcome was observed in simulations with *m* = 10 and *m* = 20, 51.6% and 68.3% of the time, respectively, and 69.0% and 48.4% in simulations with low and high ILS, respectively. This trend demonstrates that InPhyNet is sensitive to error in the input networks, with the dependence on both the number of taxa and the value of *m* further strengthening this conclusion: larger values of *m* and smaller numbers of taxa enable InPhyNet to rely more heavily on the input networks and less on the pairwise dissimilarity matrices.

This makes the decision of *m* difficult in practice, though, as higher values of *m* drastically increase runtime, but also tend to improve inference accuracy. The most important factor in determining output error, though, is input error, which does not depend on *m*. Thus, when utilizing hill-climbing algorithms that leverage the results of multiple runs such as SNaQ, PhyloNet-ML or PhyloNet-MPL, we recommend choosing smaller values of *m* and instead investing more computational resources into increasing the number of independent runs performed by the optimization software. We expect that this will provide the best tradeoff between method accuracy and total runtime.

Surprisingly, little to no difference in output accuracy was observed between PhyloNet-ML, PhyloNet-MPL, and SNaQ. Differences in runtimes between the three methods were drastic, though, indicating that the fastest method PhyloNet-MPL should be preferred. It is worth noting that these simulations were performed with version 1.0 of the Julia package `SNaQ.jl`, whose runtime has since been improved dramatically in version 1.1, so this result may no longer hold (Kolbow et al., 2026).

The results in Figure 7 show a consistent positive relationship between the amount of input data used (number of gene trees and number of base pairs in this case) and output accuracy. This trend reflects the statistical consistency results outlined above and highlights the robustness of InPhyNet as well as each of the underlying inference methods.

Our simulations utilized the AGID metric to compute a pairwise dissimilarity matrix. We show in Appendix 1 that, for generic level-1 networks, as the number of input gene trees goes to infinity, this measure does not converge on a dissimilarity matrix *D* that constructs a displayed tree of a given network as in Theorem 1. It is unclear whether other metrics may be statistically consistent in this sense or whether they would generally yield more favorable results. Future work could investigate the role that this metric plays in the overall accuracy of InPhyNet.

Additionally, one limitation of InPhyNet is that reticulations cannot be inferred between the disjoint subsets of taxa that are selected at the beginning of the inference framework. This can be accounted for by utilizing tree of blob analyses with methods like TINNiK (Allman et al., 2024) or TREE-QMC (Han and Molloy, 2025) during the subset decomposition phase. Future work could alleviate this issue by relaxing the requirement for strictly disjoint subsets of taxa, or by testing for the presence of hybridizations as the network is being built.

### Phylogeny of Green Plants

Green plants are a large, complicated group comprised of many important clades, and their analysis warrants extreme care. Our goal in this empirical analysis is not to provide novel empirical evidence on the phylogeny of green plants, but instead to illustrate how InPhyNet can be integrated into an empirical workflow to synthesize information in areas of the phylogeny that exhibit discordance.

We re-analyzed the 1,158-taxon green plant phylogeny from the One Thousand Plant Transcriptomes Initiative (1KP), focusing on regions with strong quartet concordance factor discordance that are poorly modeled by treelike phylogenies. We used InPhyNet to incorporate reticulate hypotheses where quartet-based signals conflicted, particularly among gymnosperms and ferns, into the broader phylogeny of green plants. Outside these groups, there was relatively little observed discordance, and so we treated the subtree of the 1KP inferred species tree containing these taxa as a single constraint network. We performed subset decomposition on the gymnosperms and ferns manually, utilizing past research identifying areas of discordance and the trees of blobs inferred by TINNiK (Allman et al., 2024) as guides. The resulting semi-directed network generally agrees with prior large-scale phylogenies, and provides a single, coherent hypothesis for the full reticulate phylogeny of green plants under the multispecies network coalescent model.

Within gymnosperms, the recovered network yielded a reticulate event ancestral to Gnetales that partially supports both the Gnetifer (Gnetales sister to conifers) and Gnepine (Gnetales sister to Pinaceae) hypotheses. This placement requires further analyses to validate, especially due to the systematic error in similar analyses that has been reported by Zhong et al. (2011). The reticulate event depicted in Pinaceae comprises taxa characterized by high levels of allopolyploidy. It is unclear whether SNaQ would infer a hybridization event in the presence of allopolyploidy, but PhyloNet’s maximum parsimony method is capable of inferring hybridizations in such cases (Yan et al., 2022). Whether this reticulation resolves the discordance induced by allopolyploidy or discordance induced by another source entirely requires further examination.

Within ferns, reticulate events were only identified within Polypodiidae, each of which is well-founded by prior research. The discordance among quartet concordance factors observed among ferns by One Thousand Plant Transcriptomes Initiative (2019) was primarily between the three clades depicted in Figure 9 rather than within each of the clades, so more research is warranted to further resolve this portion of the phylogeny.

While the results presented here require further research to corroborate, this analysis, particularly with respect to gymnosperms, exhibits the distinct advantage of being able to interpret reticulate events that span large portions of the phylogeny with the full context of the entire phylogeny. By preserving a well-supported backbone and targeting regions of known discordance, InPhyNet provides a coherent picture of the evolutionary history of all green plants that explicitly accounts for instances of reticulate evolution.

## Conclusions

We presented a novel method, InPhyNet, for merging constraint networks inferred on disjoint subsets of taxa. We established that InPhyNet yields correct results when the input data are correct and the disjoint subsets are chosen appropriately (see Theorem 2). Furthermore, we demonstrated that the InPhyNet algorithm is statistically consistent with respect to the amount of input data provided given that its input data are inferred in a statistically consistent manner.

Through an extensive simulation study, we verified the practicality of these theoretical findings. Key results from our simulations indicate that the accuracy of InPhyNet is highly correlated with the accuracy of its input data, and that accuracy consistently improves as the amount of input data increases. The maximum constraint network size parameter *m* had no noticeable effect on accuracy but does have a direct negative impact on runtime. The most accurate results were observed with PhyloNet-ML, though SNaQ and PhyloNet-MPL yielded similarly accurate results across the simulation study.

For a fixed value of *m*, total runtime scaled linearly with respect to the number of tips in the true species network. These results demonstrate that a pipeline utilizing accurate network inference methods, the AGID metric, and InPhyNet can be used to accurately infer networks with hundreds of tips in a very feasible time frame.

Finally, we applied InPhyNet to expand the phylogeny of green plants, successfully identifying instances of reticulate evolution in regions previously poorly supported by strictly treelike phylogenies. Collectively, these results underscore the utility of InPhyNet for robust and efficient phylogenetic network inference in complex evolutionary scenarios.

## Acknowledgements

This work was supported by the National Science Foundation (DEB-2144367 to CSL). The authors thank Jiayang Wang for his review of the proofs presented herein, Erin Molloy and Yunheng Han for their help in improving the simulation study, George Tiley for his thoughts on the empirical work, and two anonymous reviewers for their highly valuable feedback.

## Data Availability

All the original data necessary to reproduce the analyses reported in this study can be accessed through the Dryad repository available at https://doi.org/10.5061/dryad.h44j0zq0b. The InPhyNet Julia package is available via the official Julia package registry, or on GitHub at https://github.com/NathanKolbow/InPhyNet.jl. Simulation and analysis scripts are available at https://github.com/NathanKolbow/InPhyNetSimulations.

## Supplementary Material

Supplementary files, consisting of appendices detailing statistical consistency of the AGID dissimilarity measure, further details of the performance study, formal algorithms, a worked example with three constraint networks, a merge conflict example, and further details on the empirical analysis of green plants are available at SYSBIO online.

## APPENDIX 1

### Statistical Consistency OF THE AGID Metric

#### DEFINITION 8

The **internode distance** *d*_*T*_ (*u, v*) between two leaves *u* and *v* in a phylogenetic tree *T* is the sum of the lengths of the edges along the shortest path connecting *u* and *v* in *T* . The **average gene tree internode distance** (AGID) between two leaves *u* and *v* in a set of gene trees *G* is the average of the internode distances between *u* and *v* taken across all trees in *G*, namely 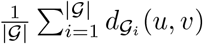.

#### THEOREM 3.

*The expected internode distance between two taxa A and B in the true gene tree *T* emerging from species tree *T* under the MSC is t*_*AB*_ + 2 *where t*_*AB*_ *is the sum of the lengths of the edges between A and B in coalescent units in the species tree *T** .

*Proof*. Under the MSC, two taxa *A* and *B* cannot coalesce until their lineages meet in the species tree **T** . The time this takes to happen is exactly the distance by which *A* and *B* are separated in **T** which we call *t*_*AB*_. Once *A* and *B* are able to coalesce, under the MSC the time until their coalescence *X* is modeled by *X ∼* Exponential(1). In the time before *A* and *B* coalesce, *both* of their branches are increasing in length, thus the total internode distance between *A* and *B* is *t*_*AB*_ + 2*X. E*(*X*) = 1, so *E*(*d*_*T*_ (*A, B*)) = *t*_*AB*_ + 2. □

#### COROLLARY 2

The expected AGID between two taxa *A* and *B* across a set of *n* true gene trees *G* emerging from species tree *T* under the MSC is *t*_*AB*_ + 2.

*Proof*. This result follows from Theorem 3 and basic properties of expectations.

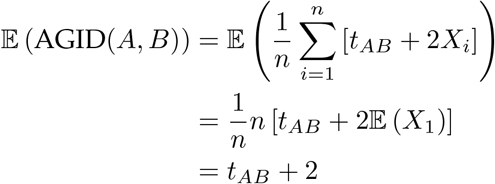

#### THEOREM 4.

*For a set of n gene trees*g**_*n*_ *inferred in an unbiased manner, the AGID metric is statistically consistent for the true species tree *T* in the sense of Definition 7 if all internal branches in *T* have non-zero length*.

*Proof*. By assuming an unbiased method of inference we have that *E*(*d*_**g**_ (*A, B*)) = *t*_*AB*_ + 2 where *t*_*AB*_ is the sum of lengths of the edges in the unique path connecting *A* and *B* in **T** .

Without loss of generality, consider taxa 1, 2 which are true neighbors in **T**and an arbitrary pair of taxa *i, j* in **T**where each of 1, 2, *i, j* are unique. The true unrooted quartet containing 1, 2, *i, j* that is displayed in must be 12| *ij* with branch lengths shown in Figure S1.1 because 1 and 2 are assumed to be true neighbors.

**Fig. S1.1.**
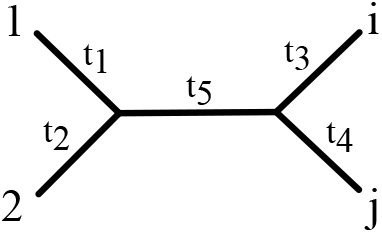
Treelike unrooted quartet and its associated edge lengths. Taxa 1 and 2 are true neighbors whereas taxa *i* and *j* are arbitrary.

It follows then that *t*_12_ = *t*_1_ + *t*_2_, *t*_*ij*_ = *t*_3_ + *t*_4_, *t*_1*i*_ = *t*_1_ + *t*_5_ + *t*_3_, and so on. Thus, E(*Q*_*k*_^*(T)*^(*M_D*(*G*_*n*_); 1, 2, *i, j*)) = −2*t*_5_ for *k* = 1, 2 where *M_D*(*G*_*n*_) = *d*_{*G*_*n*_}, so the AGID metric is statistically consistent for *T* by Definition 7.

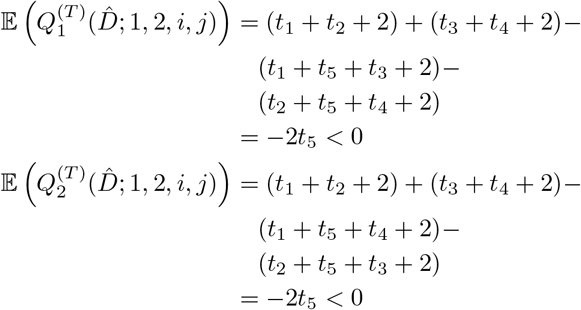

#### Quarnet Derivations

For a quarnet *N* with taxa labeled 1, 2, *A, B* as in Figures S1.2 to S1.7, we write 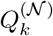 (or simply *Q*_*k*_ when *N* is unambiguous) as shorthand for 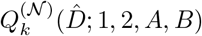, where 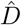 is the AGID matrix. We now derive 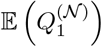 and 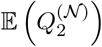 for each possible level-1, binary, semi-directed quarnet *N*. To do so, we derive each 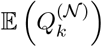 conditioned on each possible coalescent history of *N* before applying the Law of Total Probability. The derivations of the probabilities of each coalescent history for quarnets 1, 3, 4, 5, and 6 are outlined in Section 1 of the Supporting Information of Solís-Lemus and Ané (2016), and we additionally include quarnet 2 here for completeness.

Note that for each quarnet there are at most 8 possible labelings that preserve the neighborhood of taxa 1 and 2. These alternative labelings can be obtained by applying combinations of the following moves: (a) swap the labels for taxa 1 and 2, (b) swap the labels for taxa *A* and *B*, or (c) swap the labels for taxa *A* and *i* and swap the labels for taxa *B* and *j*. Notice that moves (a) and (b) simply adjust the labels such that *Q*_1_ becomes exactly *Q*_2_ and *Q*_2_ becomes exactly *Q*_1_, and move (c) simply reorders the terms in *Q*_1_ and *Q*_2_. Thus, *Q*_1_ and *Q*_2_ are invariant to the labeling of a given quarnet, so we derive their expectations for a single labeling for each quarnet.

Let 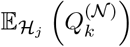 denote the expected value of 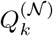 for quarnet *N* under coalescent history *ℋ*_*j*_. Then, the results for each quarnet are as follows.

**Fig. S1.2.**
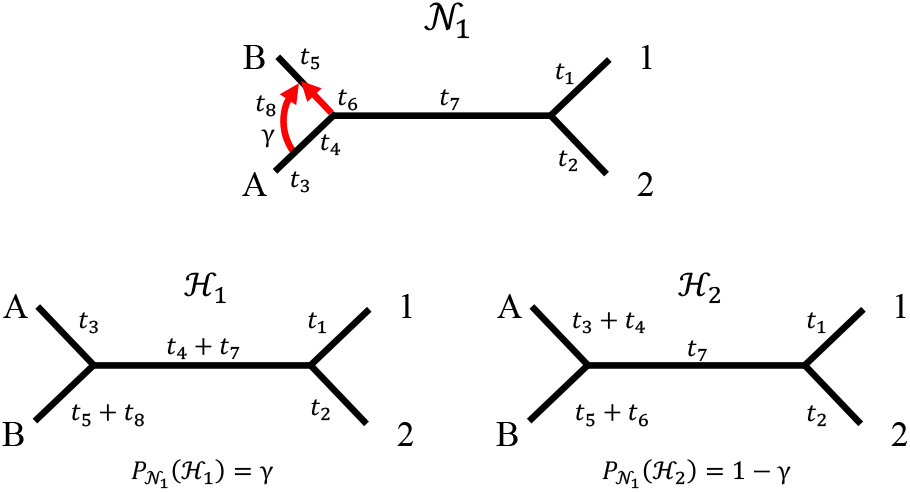
Quarnet 1, its coalescent histories *ℋ*_*j*_, and their associated probabilities.

**Fig. S1.3.**
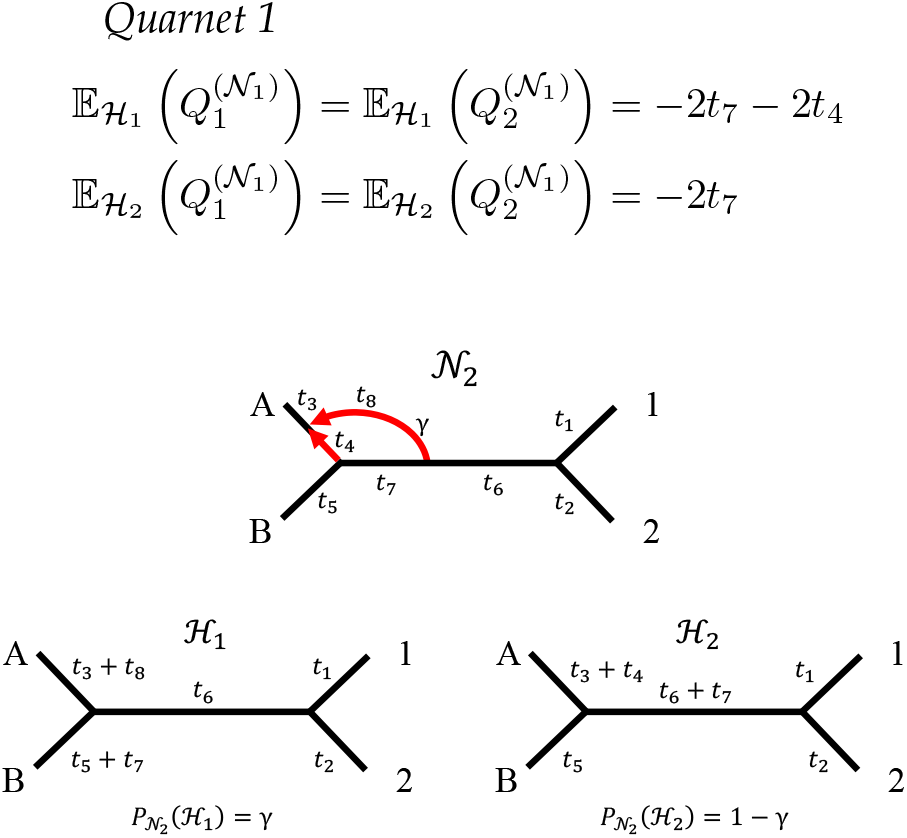
Quarnet 2, its coalescent histories *ℋ*_*j*_, and their associated probabilities.

**Fig. S1.4.**
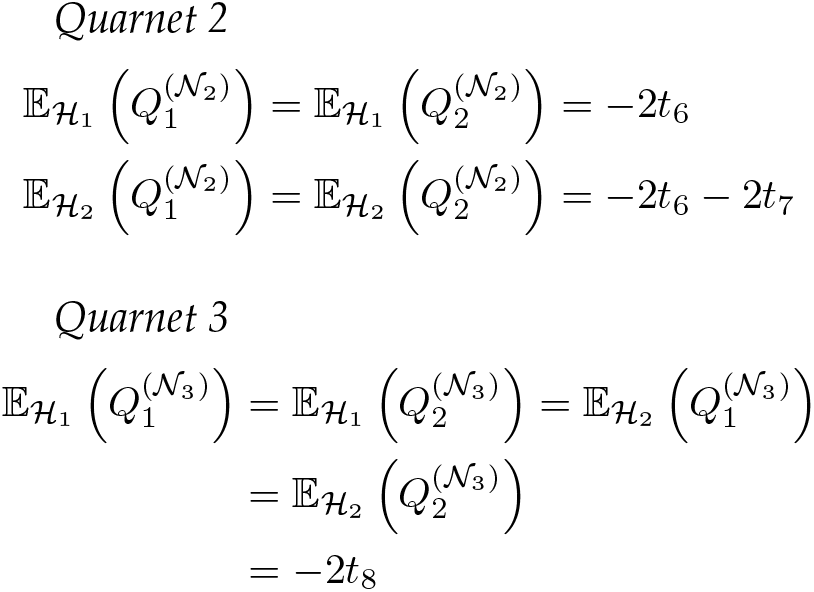
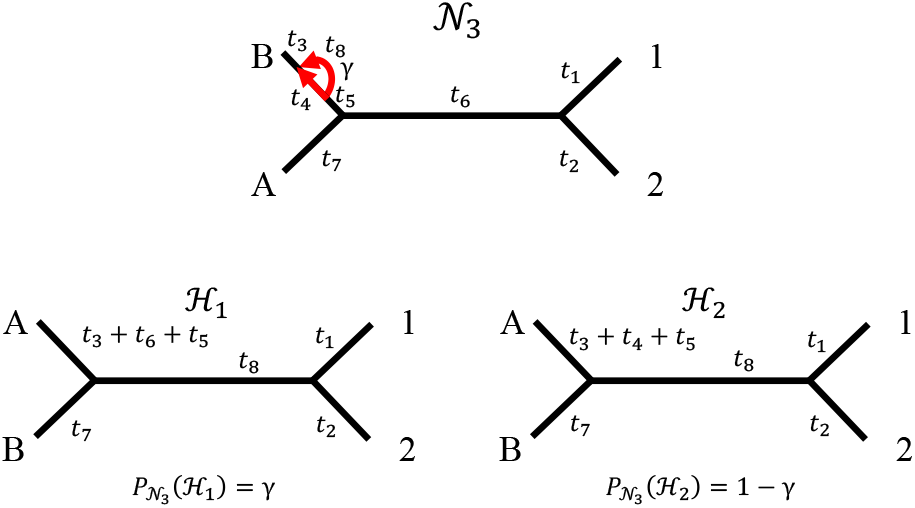
Quarnet 3, its coalescent histories *ℋ*_*j*_, and their associated probabilities

**Fig. S1.5.**
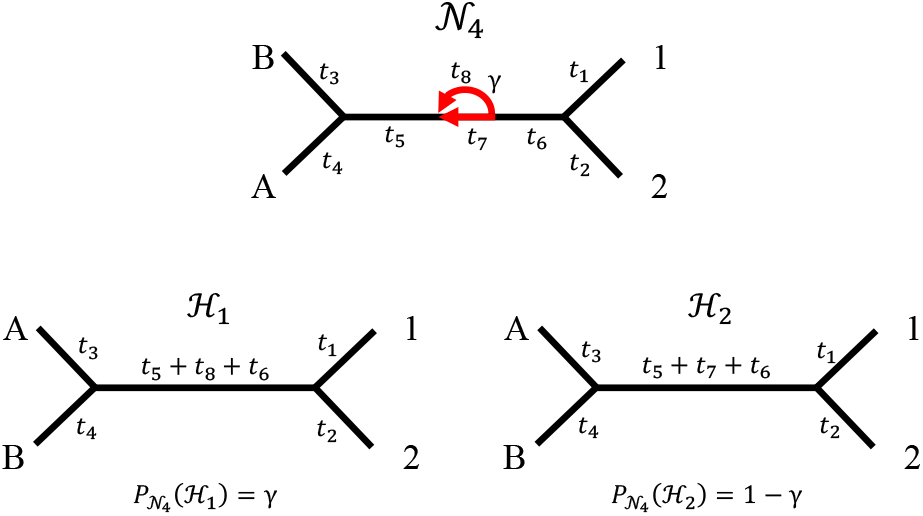
Quarnet 4, its coalescent histories *ℋ*_*j*_, and their associated probabilities

**Fig. S1.6.**
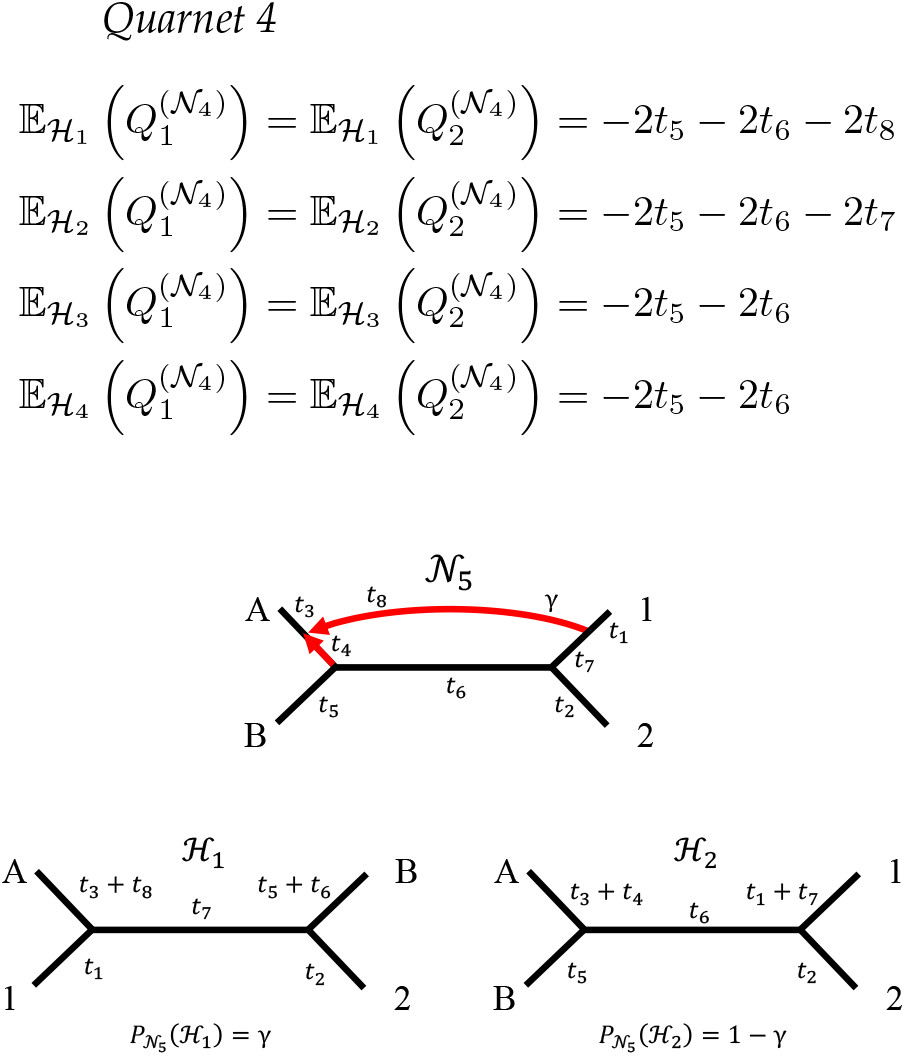
Quarnet 5, its coalescent histories *ℋ*_*j*_, and their associated probabilities.

**Fig. S1.7.**
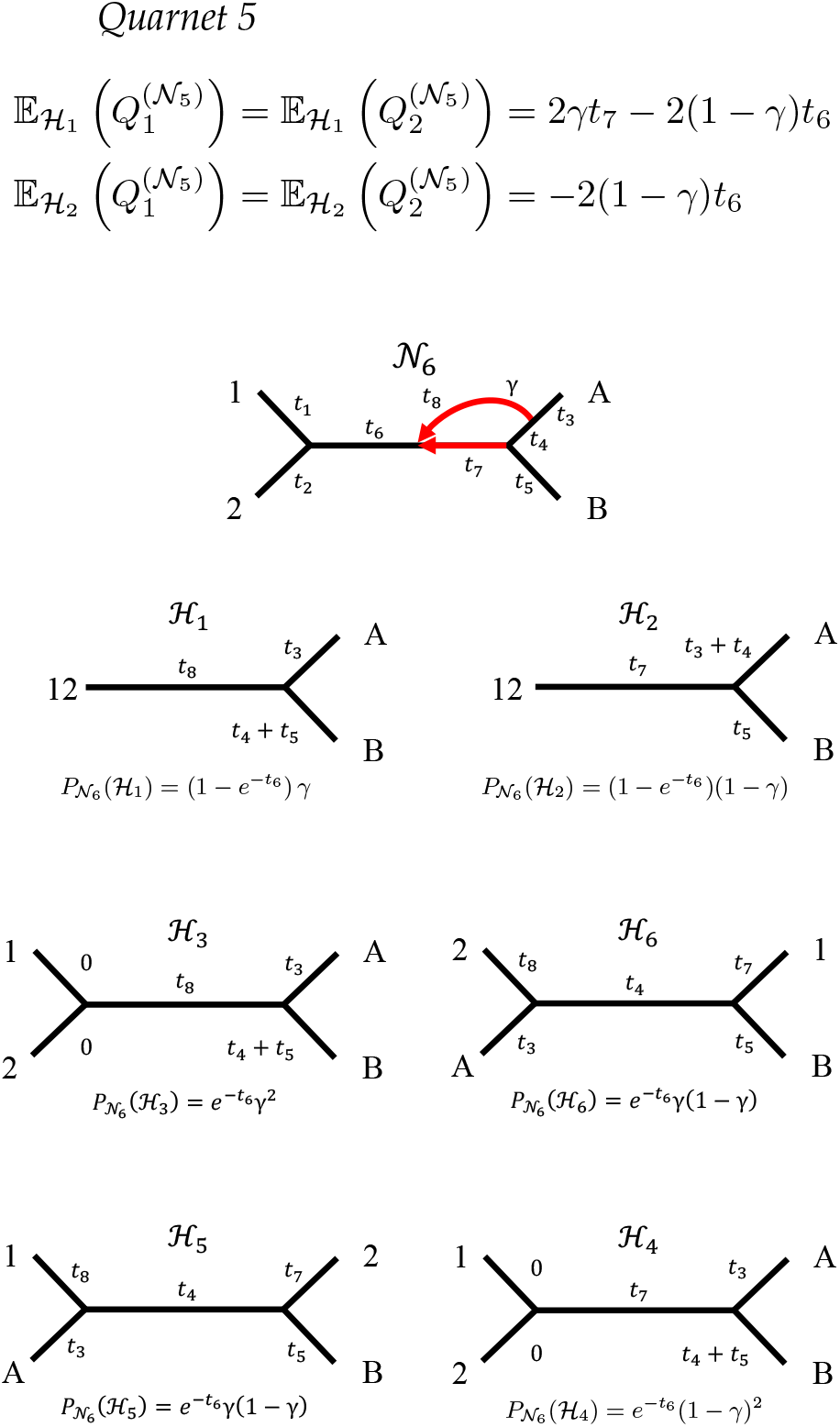
Quarnet 6, its coalescent historie *ℋ*_*j*_, and their associated probabilities.

#### Quarnet 6

Under coalescent histories _1_ and _2_, taxa 1 and 2 are assumed to have merged prior to reaching the hybrid node in _6_, so E(*D*_12_) takes on the following new form:

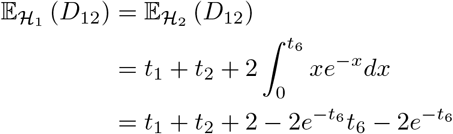

With this result, we compute each 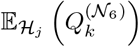.

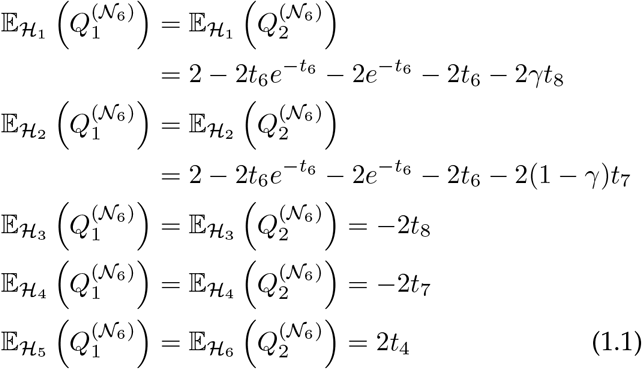

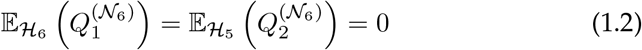

Note the differing pattern in the indices of Equations 1.1 and 1.2.

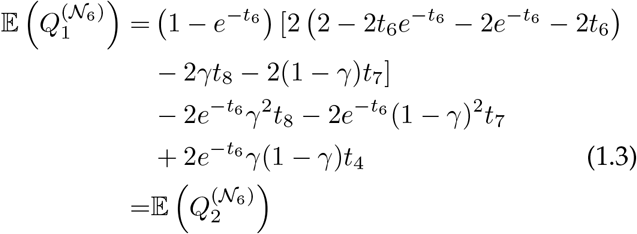

##### Theorem 5.

*For a set of n gene trees, the AGID metric is* ***not*** *statistically consistent for an arbitrary true species network N in the sense of Definition 7*.

*Proof*. Choose *N* t o be *N*_6_ with *t*_4_ = 1, *t*_6_ = *t*_7_ = *t*_8_ = 0, *γ* = 0.5. Then, 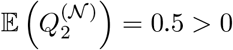.

Quarnets *N*_5_ and *N*_6_ provide clear counter-examples to the property of statistical consistency, but it is unclear how salient of an impact these results would have in practice. For 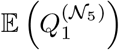 to be non-negative, which would lead to AGID not being statistically consistent for some displayed tree of the true species network, we need *γt*_7_ *⩾* (1 *γ*)*t*_6_ in *N*_5_. This condition seems particularly lenient, but can be largely avoided by including samples from two individuals of each taxon or species. In this case, the labeling of *N*_5_ with 1 or 2 under the hybrid node would no longer be possible, because any leaf under the hybrid node would have a true neighbor that is also under the hybrid node. Then, only the labeling as shown in Figure S1.6 is possible. To satisfy the given inequality in this case, true neighbors 1 and 2 would need to be particularly far from one another, leading to a large *t*_7_, while their most recent common ancestor would need to be very close to the most recent common ancestor of *i* and *j*, leading to a small *t*_6_. Whether such a case is particularly common or plausible is unclear, and would require more research.

In the case of 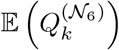, *k* = 1, 2, a more complicated relationship arises (see Equation 1.3). In this case, what is required for AGID to *not* be statistically consistent for some displayed tree of the true species network is that, in general, *γ* must not be close to 0 or 1, *t*_6_, *t*_7_, and *t*_8_ must be very small relative to *t*_4_, *or t*_4_ must be very, very large. This reflects a situation where there is moderate to high levels of genetic exchange at the hybrid node while the parents of the hybrid node in *N*_6_ are very far from one another relative to the distances represented by *t*_6_, *t*_7_, and *t*_8_. These conditions are highly reminiscent, though not identical, to the conditions necessary for a network to be anomalous (see Ané et al., 2024).

## APPENDIX 2

### Performance Study Further Details

#### Statistics on Generated Data

For each simulation, we computed the gene tree estimation error (GTEE) as the average hardwired cluster dissimilarity (HWCD) between each true gene tree and its associated estimated gene tree, divided by the maximum possible HWCD value (2 |*X*| *−*6 where |*X*| is the number of taxa in the tree). This value varied heavily across simulation settings, the distribution of which is shown in Figure S2.1.

**Fig. S2.1.**
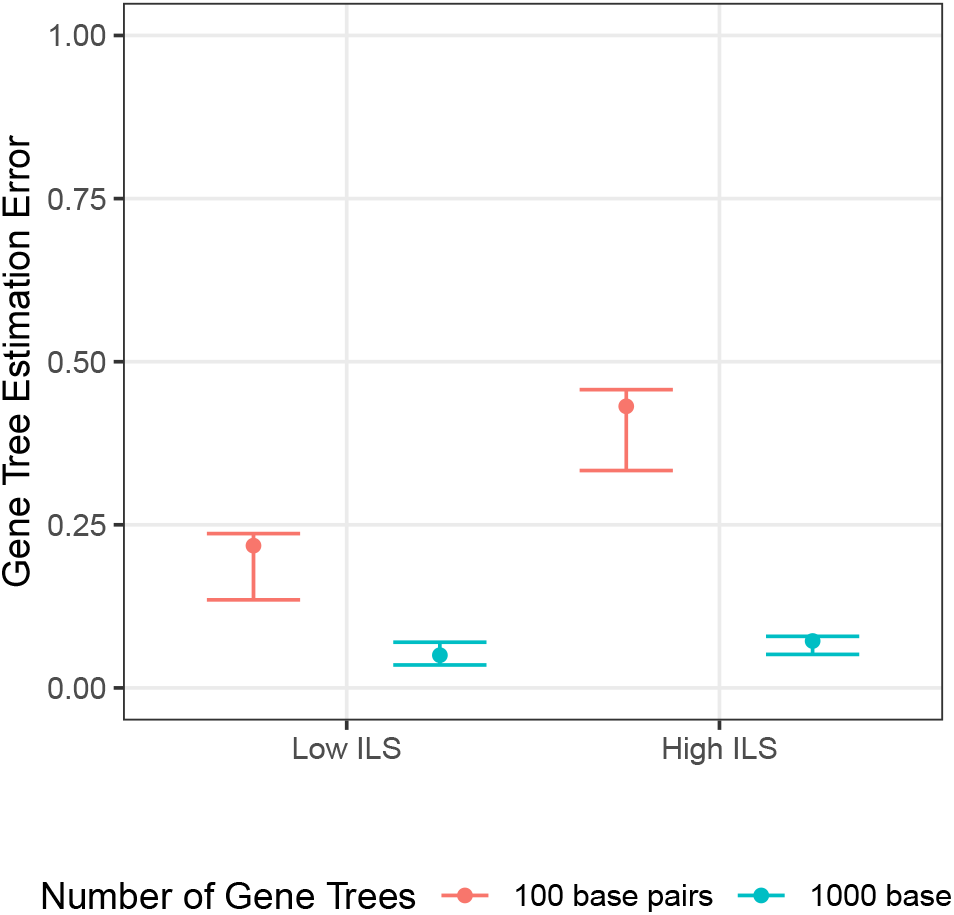
Distribution of gene tree estimation errors (GTEEs) across various simulation settings. Values did not vary significantly across simulations with differing number of taxa or gene trees. Values are irrelevant to simulations that utilized Squirrel because Squirrel utilizes MSAs as input, not estimated gene trees.

**Table 2.1.** Average gene tree estimation error (GTEE) for each combination of number of taxa, incomplete lineage sorting (ILS), and number of base pairs (NBP) in the MSA.

| NBP | ILS | Number of Taxa | Avg. GTEE |
| --- | --- | --- | --- |
| 100 | Low | 25 | 0.221 |
| 100 | Low | 50 | 0.233 |
| 100 | Low | 100 | 0.233 |
| 100 | Low | 200 | 0.267 |
| 100 | High | 25 | 0.376 |
| 100 | High | 50 | 0.419 |
| 100 | High | 100 | 0.449 |
| 100 | High | 200 | 0.464 |
| 1000 | Low | 25 | 0.082 |
| 1000 | Low | 50 | 0.102 |
| 1000 | Low | 100 | 0.147 |
| 1000 | Low | 200 | 0.152 |
| 1000 | High | 25 | 0.102 |
| 1000 | High | 50 | 0.094 |
| 1000 | High | 100 | 0.140 |
| 1000 | High | 200 | 0.105 |

#### Simulation Software Parameters

Default parameters were used for IQ-TREE, ASTRAL v5.7.1, and Squirrel. For IQ-TREE, this means that IQTREE’s ModelFinder functionality was used and the true underlying model of evolution was never specified when inferring gene trees. Specific divergences from default parameters in other software are enumerated below. Any parameters that are unmentioned were left at their default values.

##### seq-gen

For each true gene tree, one MSA was generated with a single sample for each tip in the gene tree. Base pairs were simulated under the F81 model of evolution (Felsenstein, 1981) with state frequencies *π*_*A*_ = *π*_*T*_ = 0.3 and *π*_*C*_ = *π*_*G*_ = 0.2. The branch length scaling parameter (`-s` ) was set to 0.05.

##### TINNiK

Parameters *α* = 0.01 and *β* = 0.99 were used.

##### SNaQ

The first estimated gene tree was used as the starting topology for each independent run of SNaQ. Additionally, 20 independent runs were performed for each constraint network. For a given subset of taxa, the maximum number of hybrids parameter (`hmax` ) was chosen to be the number of hybrids in the subnetwork of the true species network spanned by that subset of taxa.

##### PhyloNet

As with SNaQ, the maximum number of hybrids in each constraint network was chosen to be the number of hybrids in the subnetwork of the true species network spanned by that constraint network’s subset of taxa. Ten independent runs were performed for each constraint network, and each round of inference was performed with 5 processors.

## APPENDIX 3

### Algorithms

Below, we provide formal algorithms for each step of InPhyNet, along with informal proofs of runtime complexity upper bounds. The following definition is used in Algorithms 3 and 4 below.

#### Definition 9

Denote the set of all paths from *u* to *w* in constraint network *C*_*k*_ as 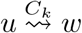. The path 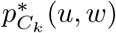 (or *p*^*\**^when *C*_*k*_ is unambiguous) is the shortest path belonging to 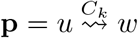, unless **p** contains a path of length 4 and a path of length 5. Then, 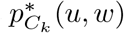 is the path of length 5. Here, as in Definition 6, the length of a path is its number of vertices. This exception is in place specifically for case 5 in Figure 2 in the main text

For the convenience of the reader, Table 3.1 offers descriptions of variables that are used frequently throughout the following algorithms.

**Table 3.1.**
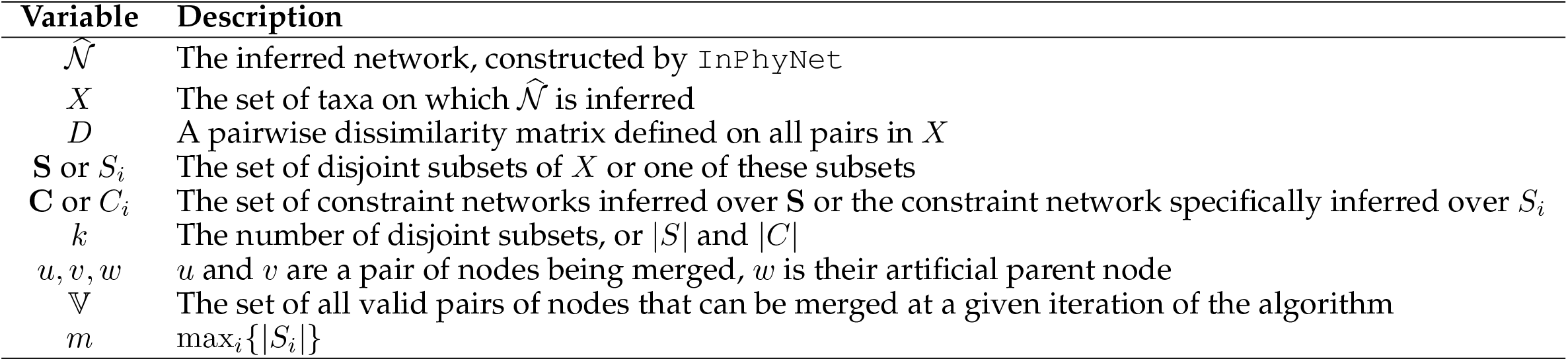
Descriptions of variables used frequently throughout the algorithms below.

#### Algorithm 1

InPhyNet(*X, D*, **C**)

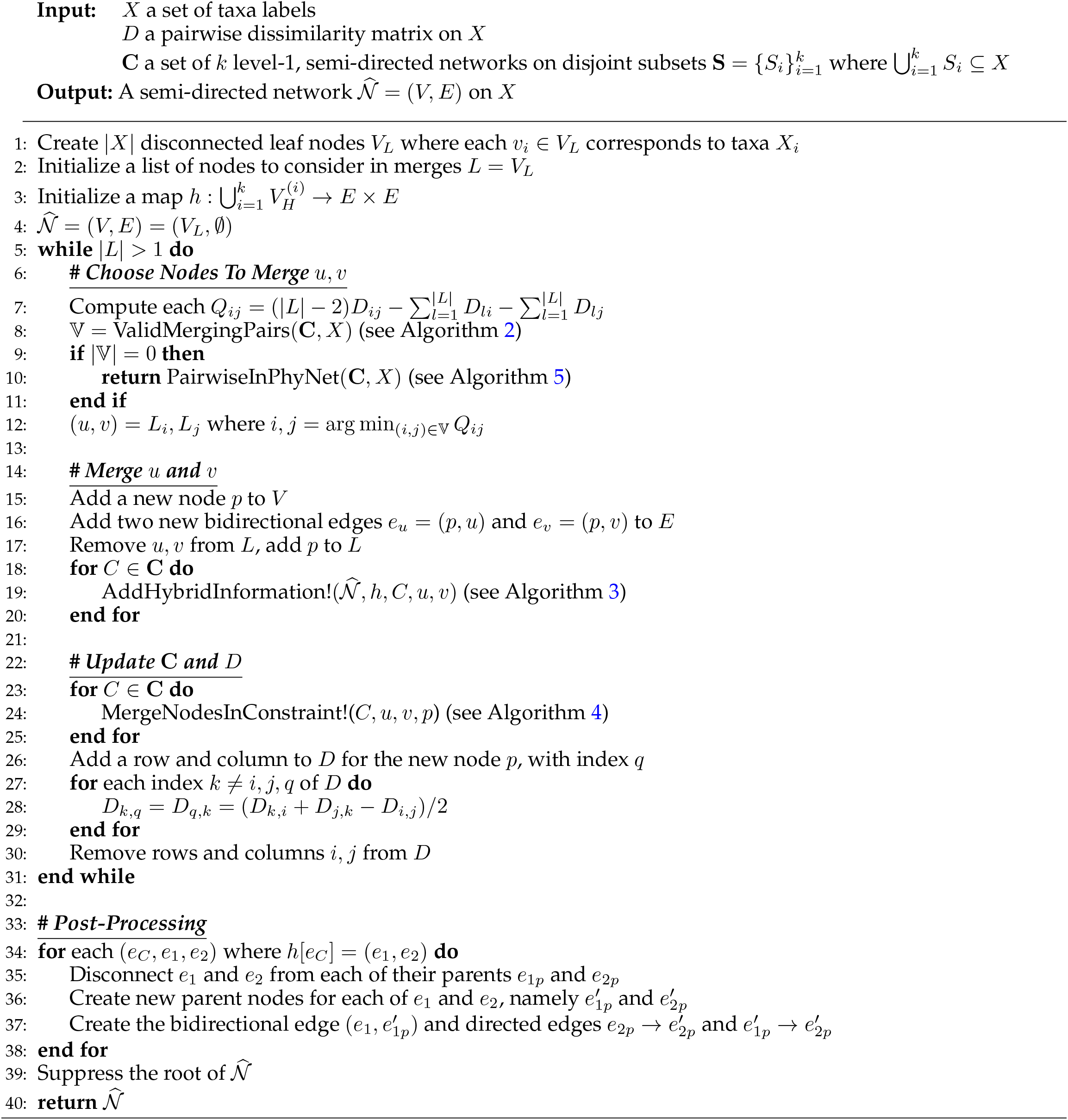

In the following, assume that all subsets are of equal size, or take *m* to be the size of the largest subset. In the ***Choose Nodes To Merge*** *u, v* section, there are at most on the order of *mk* valid merging pairs in *V* where *k* is the total number of disjoint subsets (see Algorithm 2), so we need only calculate each *Q*_*ij*_ for these pairs. This leaves us with a runtime complexity of *O*(*k*^2^*m*^2^). Then, the first 3 lines of ***Merge*** *u* ***and*** *v* happen in constant time while the for loop is executed *k* times, and its contents have runtime complexity *O*(*m* + *r*) (see Algorithm 3), leaving it with runtime complexity *O*(*km* + *kr*). Finally, examining each for loop in the section ***Update* C *and*** *D* gives the runtime complexity *O*(*km* + *kr*)) (see Algorithm 4). Each of these steps happens |*X*| times, so the overall runtime complexity, excluding post-processing, given that `PairwiseInPhyNet` is never called, is *O*(|*X*|*k*^2^*m*^2^ + |*X*|*kr*). Notice that, by definition, 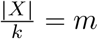, so this complexity reduces to *O*(|*X*|^3^ + |*X*|*kr*).

In the worst case, this algorithm is executed until the final iteration where *V* = 0 on line 9 leads to `PairwiseInPhyNet` being called. The runtime complexity of `PairwiseInPhyNet` is *k* times the runtime complexity of this algorithm (see Algorithm 5; `PairwiseInPhyNet` is never called when `InPhyNet` is called from `PairwiseInPhyNet` ). When `InPhyNet` is called from inside of `PairwiseInPhyNet`, *k* is fixed as 2. Further, `InPhyNet` is called a total of *k −* 1 times from within `PairwiseInPhyNet` . Thus, the worst-case complexity of the `InPhyNet` algorithm is *O*(|*X*|*k*^2^*m*^2^ + |*X*|*kr* + (*k −* 1)(2^2^|*X*|*m*^2^ + 2|*X*|*r*)) = *O*(|*X*|3 + |*X*|*kr*). Thus, the runtime complexity of the `InPhyNet` algorithm is *O*(|*X*|3 + |*X*|*kr*) or *O*(|*X*|3 + |*X*|*r*) if *k* is treated as fixed.

#### Algorithm 2

ValidMergingPairs(**C**, *X*)

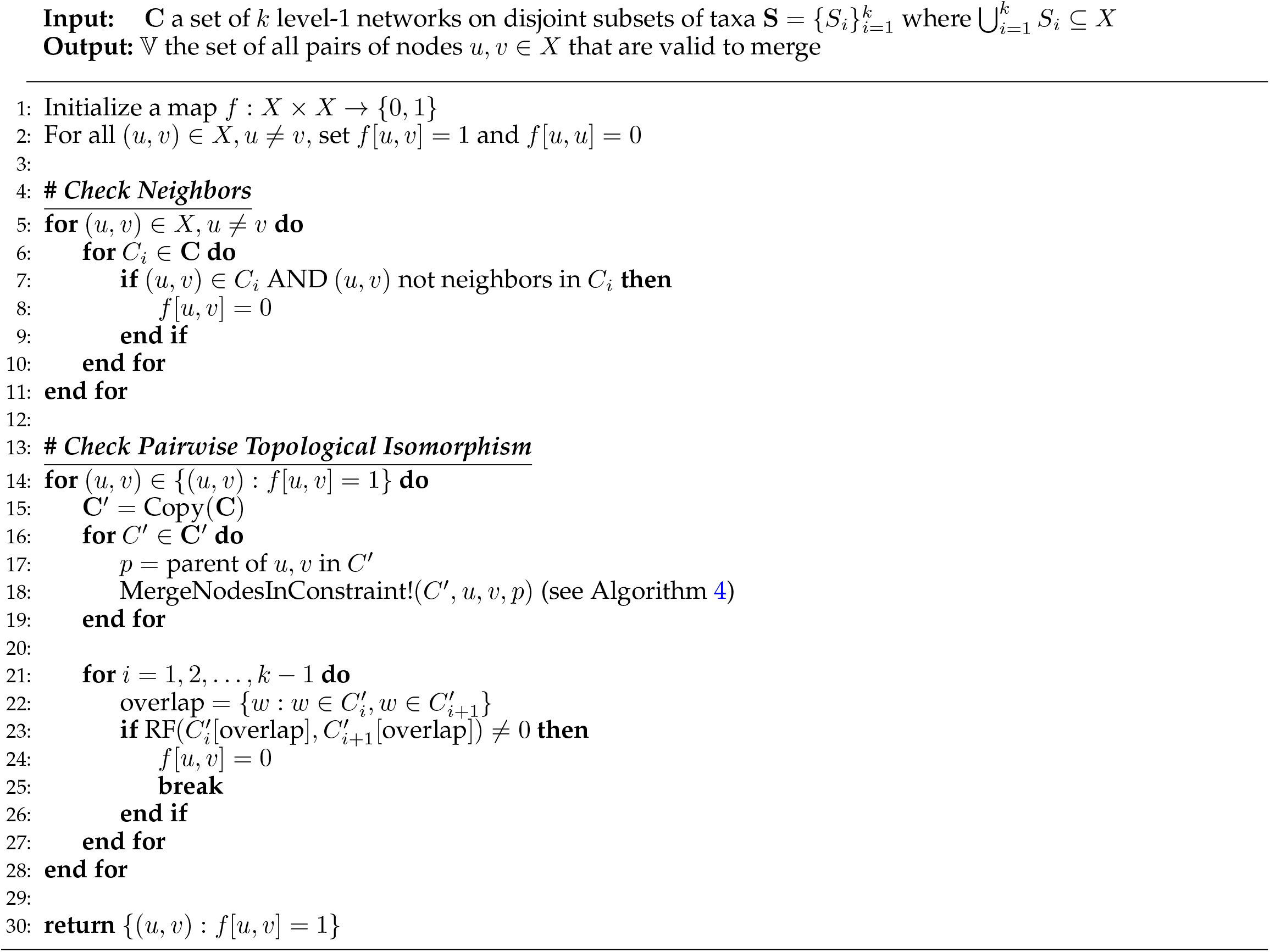

The overall runtime of this algorithm can be broken down into two sections: the portion marked ***Check Neighbors*** and the portion marked ***Check Pairwise Topological Isomorphism***. Checking neighborhood status can be done more efficiently by initially marking all pairs of taxa as non-neighbors, then iterating over each leaf in each network *C*_*i*_ ∈ **C** and checking whether that leaf has a sibling. Checking for a sibling can be done in constant time, and each network has at most *m* taxa, so this process has runtime complexity *O*(*mk*).

A given unrooted tree with *m* can have at most 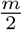 sets of neighbors (in the case where the tree is perfectly balanced), and, by our definition of neighbor, networks can have one additional neighbor per reticulation. Thus, the most neighbors that a single network can have is 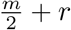. Then, Algorithm 4 has runtime complexity *O*(*m* + *r*), and the Robinson-Foulds (RF) distance metric can be computed in linear time (Day, 1985). Thus, the runtime complexity of the second portion of this algorithm is *O*(*k*(*m* + *r*)^2^ + (*m* + *r*)*km*, bringing the overall algorithm runtime complexity to *O*(*k*(*m* + *r*)^2^).

#### Algorithm 3

AddHybridInformation!(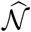, *h, C*_*i*_, *u, v*)

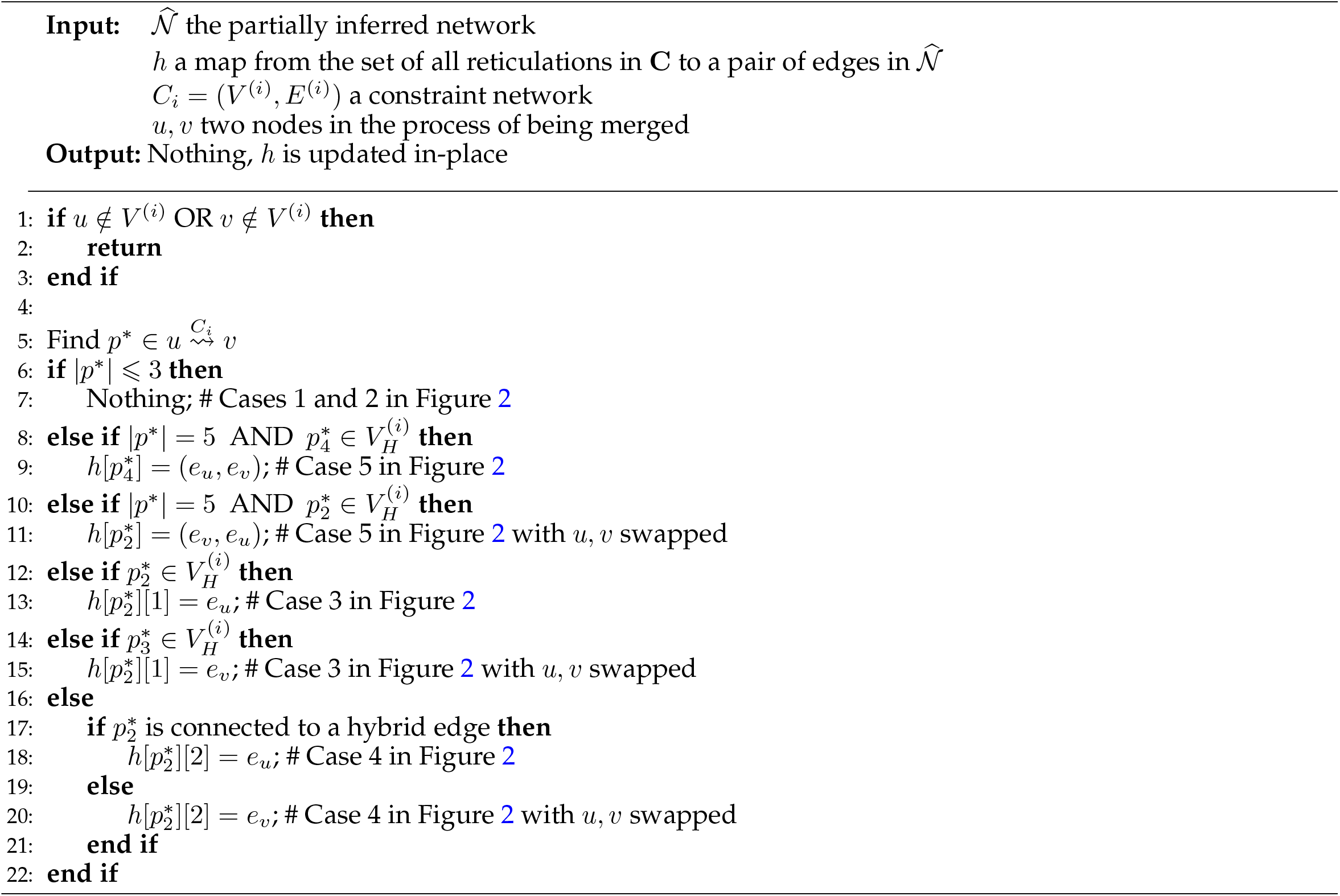

Each portion of this algorithm runs in constant time except for line 6 when 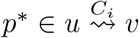 is obtained. In the worst case with a pathfinding algorithm like A^*\**^, the runtime complexity of this step is upper bounded by the number of edges in the graph.

To derive an upper bound for this value, we examine the maximum number of edges that can appear in tree-based networks, a parent class to level-1 networks (see Kong et al., 2022 for further details). All tree-based networks can be constructed by taking a tree and attaching a series of reticulations to the tree. To construct a semi-directed level-1 network, we will begin this process with an unrooted phylogenetic tree, which has 2*m −*3 total edges when *m* is the number of tips in the network. Then, each time a reticulation is added to this topology, the most new edges that can be created is three (one from splitting the edge at the origin of the reticulation, one from splitting the edge at the destination of the reticulation, and the third being the reticulation itself). Thus, each reticulation adds at most 3 new edges to the network, and we have an upper bound of 2*m* + 3*r −*3 where *m* is the number of tips in the network and *r* is the number of reticulations in the network. This gives an upper-bounded runtime to Algorithm 4 of *O*(*m* + *r*).

#### Algorithm 4

MergeNodesInConstraint!(*C*_*i*_, *u, v, w*)

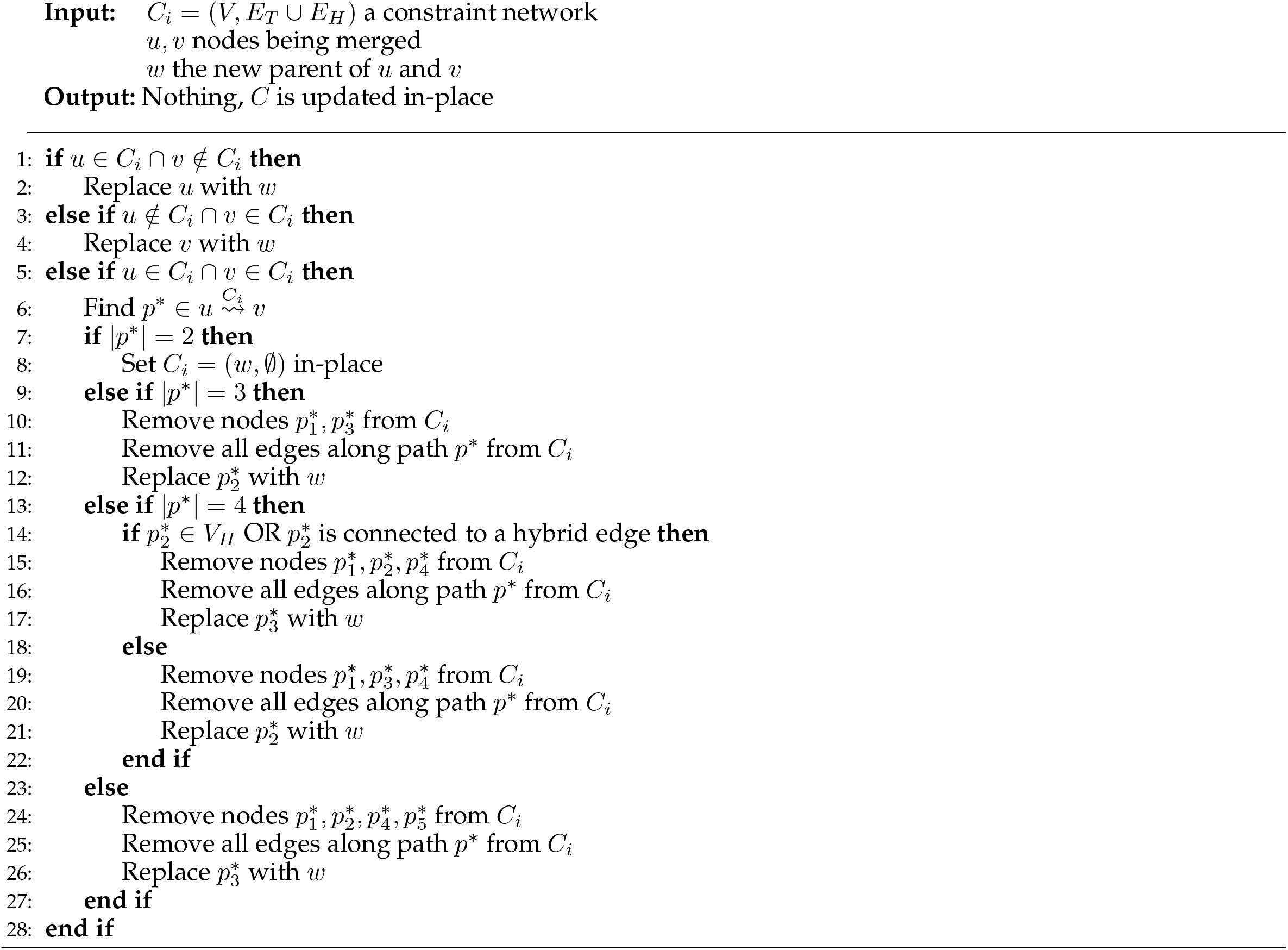

Following the same reasoning as for Algorithm 3, this algorithm’s runtime is *O*(*m* + *r*).

#### Algorithm 5

PairwiseInPhyNet(**C**, *D, X*)

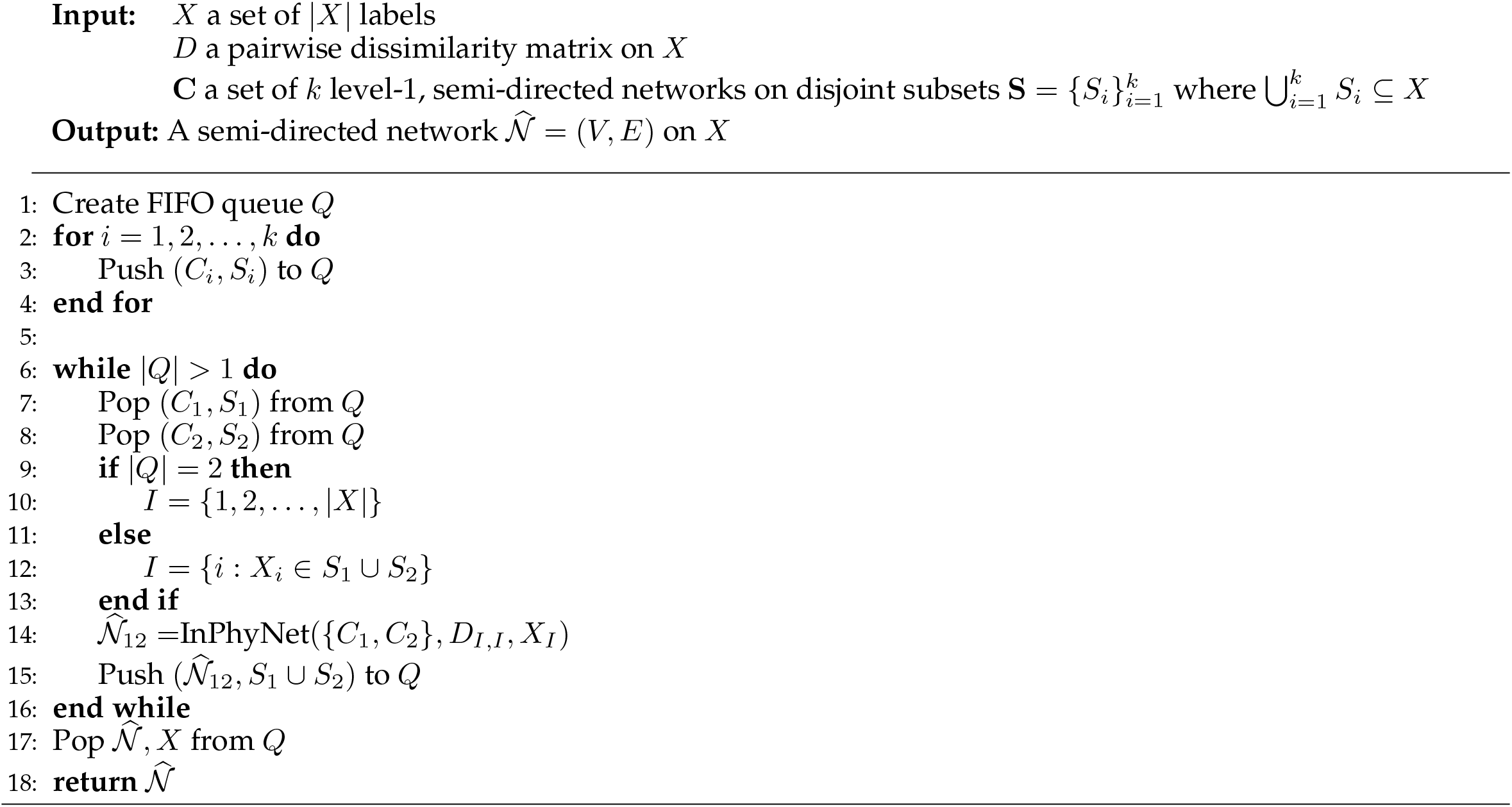

The **while** loop on line 6 will execute *k* −1 times, so the runtime complexity of this function will be *k* − 1 times the runtime complexity of the `InPhyNet` function. See the discussion under Algorithm 1 for further details.

## APPENDIX 4

### Worked Example with Three Constraints

Figure S4.1 on the following page walks through an example of the InPhyNet algorithm with three constraint networks and three reticulations. Note that while constraints and portions of the inferred network are drawn as rooted, they are in fact semi-directed. At each iteration, after two nodes are merged, their new corresponding node in each constraint is highlighted in red font. Additionally, changes to the inferred network are highlighted in red font at each iteration.

As hybrid information is added to the inferred network, edges are colored light blue and marked with (i) the name of the hybrid to which they correspond (H1 for the first constraint, H2 for the second, and H3 for the third), and (ii) the direction of the reticulation relative to the edge (in or out). This information is displayed at each iteration where it first appears, but is hidden in subsequent iterations for improved clarity. All of this information is displayed simultaneously at the end of the figure where the hybrid connections and unrooting processes are shown.

To perform this exact same analysis in Julia, perform the following in the Julia REPL:

~~~
using InPhyNet
loadworkedexample()
~~~

Note that this example displays an interesting property of InPhyNet: the constraint networks need to be level-1, but the output network does not necessarily need to be. In this case, the inferred network given by InPhyNet is level-3 with closely connected hybridization events.

**Fig. S4.1.**
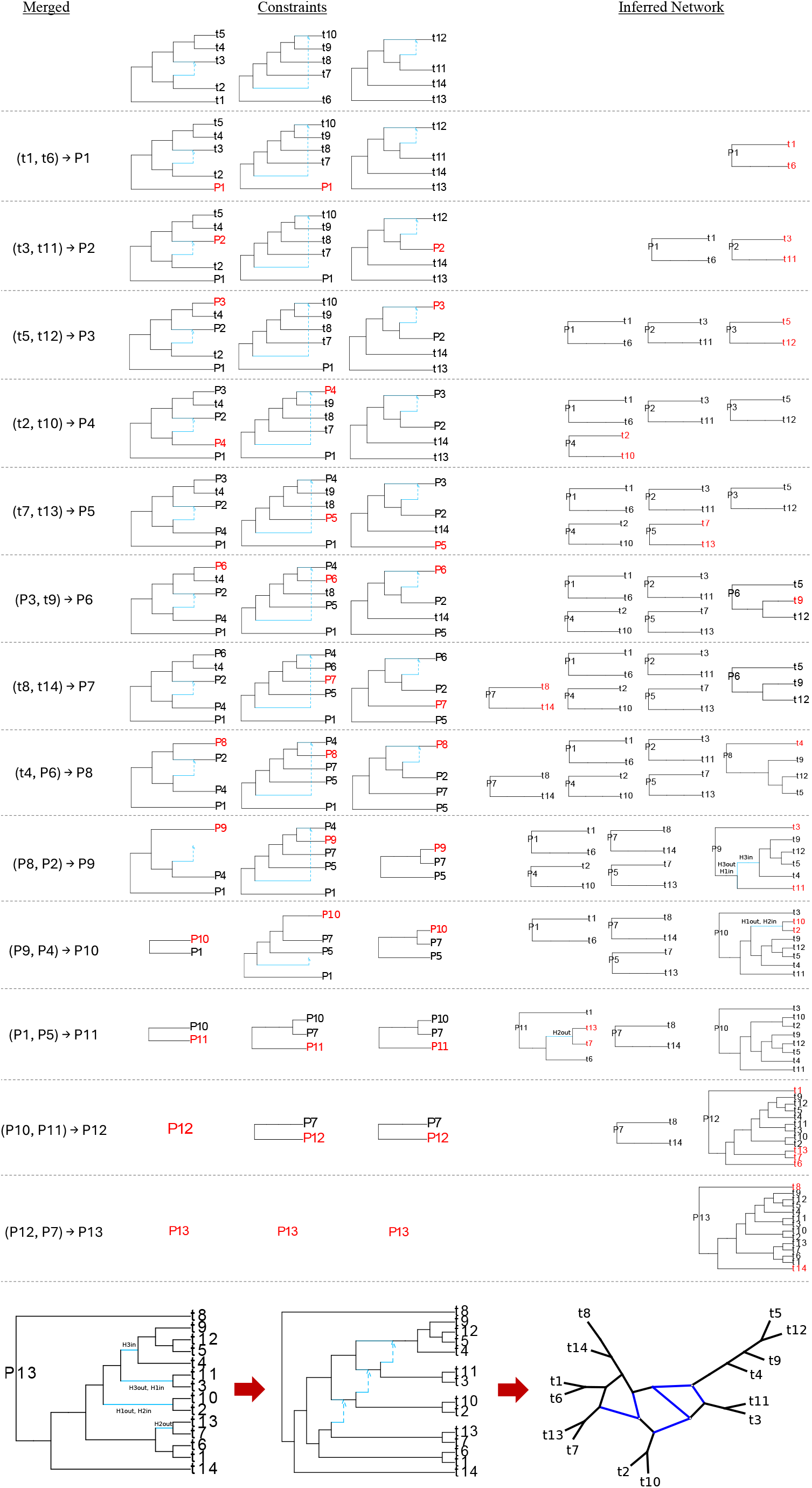

## APPENDIX 5

### Merge Conflict Example

Here, we provide an example of when the `InPhyNet` algorithm would fail without Algorithm 5. Below we provide a dissimilarity matrix *D* (Table 5.1) and a set of constraints **C** that leads to this result. Additionally, we show how the constraints change throughout the algorithm, eventually reaching a point where there do not exist any pairs of nodes that can be merged.

**Table 5.1.** Example dissimilarity matrix that leads to conflict.

| | $t_1$ | $t_2$ | $t_3$ | $t_4$ | $t_5$ | $t_6$ | $t_7$ | $t_8$ | $t_9$ | $t_{10}$ | $t_{11}$ | $t_{12}$ |
| --- | --- | --- | --- | --- | --- | --- | --- | --- | --- | --- | --- | --- |
| $t_1$ | 0.0 | 0.83 | 0.99 | 0.54 | 0.19 | 0.49 | 0.22 | 0.61 | 0.28 | 0.21 | 0.94 | 0.73 |
| $t_2$ | - | 0.0 | 0.46 | 0.86 | 0.45 | 0.47 | 0.08 | 0.17 | 0.61 | 0.94 | 0.43 | 0.74 |
| $t_3$ | - | - | 0.0 | 0.5 | 0.92 | 0.47 | 0.47 | 0.35 | 0.46 | 0.03 | 0.18 | 0.1 |
| $t_4$ | - | - | - | 0.0 | 0.5 | 0.69 | 0.07 | 0.5 | 0.31 | 0.67 | 0.67 | 0.86 |
| $t_5$ | - | - | - | - | 0.0 | 0.67 | 0.83 | 0.27 | 0.5 | 0.21 | 0.19 | 0.96 |
| $t_6$ | - | - | - | - | - | 0.0 | 0.93 | 0.98 | 0.93 | 0.28 | 0.65 | 0.17 |
| $t_7$ | - | - | - | - | - | - | 0.0 | 0.46 | 0.92 | 0.03 | 0.8 | 0.21 |
| $t_8$ | - | - | - | - | - | - | - | 0.0 | 0.46 | 0.24 | 0.1 | 0.65 |
| $t_9$ | - | - | - | - | - | - | - | - | 0.0 | 0.12 | 0.2 | 0.2 |
| $t_{10}$ | - | - | - | - | - | - | - | - | - | 0.0 | 0.65 | 0.7 |
| $t_{11}$ | - | - | - | - | - | - | - | - | - | - | 0.0 | 0.81 |
| $t_{12}$ | - | - | - | - | - | - | - | - | - | - | - | 0.0 |

We use the notation (*t*_*i*_, *t*_*j*_) *→ t*_*k*_ to indicate that nodes *t*_*i*_ and *t*_*j*_ have been chosen to be merged, with the new artificial parent that will be replacing them in the constraint networks and dissimilarity matrix denoted *t*_*k*_. Then, this dissimilarity matrix along with the constraint networks shown in the top-left of Figure S5.1 lead to a merge conflict.

**Fig. S5.1.**
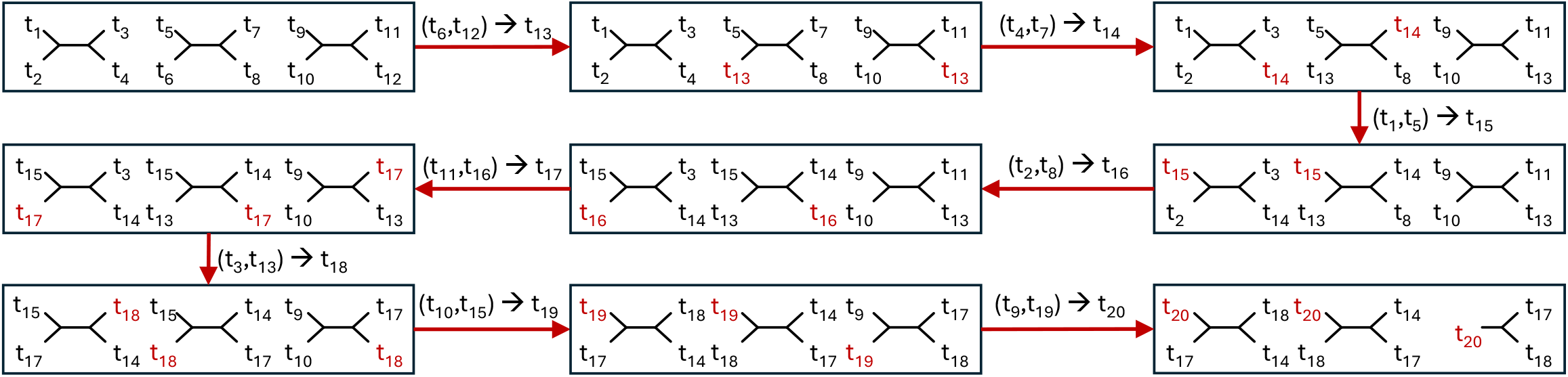
Example of a conflict that can arise in InPhyNet when more than 2 constraint networks are used as input. The set of constraint networks at each step of the algorithm is outlined in a black rectangle. Red arrows between these rectangles indicate the algorithm progressing forward one step. The label on each of these arrows (*t*_*i*_, *t*_*j*_) *→ t*_*k*_ indicates that nodes *t*_*i*_ and *t*_*j*_ are being merged and replaced by their new artificial parent *t*_*k*_. After each step of the algorithm, nodes that were modified are given red font. The algorithm begins in the top left and ends in the bottom right. At the end of the algorithm, no valid pairs of nodes exist that can be merged.

Notice at the end of the algorithm example shown in Figure S5.1 that there are 4 unique artificial taxa remaining among the constraints: *t*_14_, *t*_17_, *t*_18_, and *t*_20_. The third constraint is a triplet, so it does not restrict the set of valid nodes. The first and second constraints, on the other hand, contain the same 4 tips but with differing topologies. This means that every possible pair of these remaining 4 taxa will be valid in one of these networks, but invalid in the other.

To perform this exact same analysis in Julia, perform the following in the Julia REPL:

~~~
using InPhyNet
loadconflictexample()
~~~

## APPENDIX 6

### Green Plants Subset Decomposition

Here, we enumerate the subsets (including outliers) with which we inferred constraint networks using SNaQ (Solís-Lemus and Ané, 2016). Networks were inferred with *h* = 0, 1, 2, 3 reticulations for each subset. We provide the trees of blobs inferred with TINNiK (Allman et al., 2024) that were used to perform manual subset decomposition in Figures S6.1 and S6.3. Additionally, we provide figures displaying the model selection decisions for these subsets in Figures S6.2 and S6.4.

#### Gymnosperms

- Subset 1 (21 taxa): *Sundacarpus amarus, Prumnopitys andina, Halocarpus bidwillii, Microstrobos fitzgeraldii, Phyllocladus hypophyllus, Parasitaxus usta, Manoao colensoi, Lagarostrobos franklinii, Saxegothaea conspicua, Microcachrys tetragona, Acmopyle pancheri, Podocarpus coriaceus, Podocarpus rubens, Nageia nagi, Retrophyllum minus, Dacrycarpus compactus, Dacrydium balansae, Falcatifolium taxoides, Falcatifolium taxoides, Botryococcus braunii, Chloromonas perforata*
- Subset 2 (15 taxa): *Sciadopitys verticillata, Cephalotaxus harringtonia, Amentotaxus argotaenia, Torreya taxifolia, Torreya nucifera, Austrotaxus spicata, Pseudotaxus chienii, Taxus baccata, Taxus cuspidata, Araucaria rulei, Wollemia nobilis, Agathis robusta, Araucaria sp., Coccomyxa pringsheimii, Pleurastrum insigne*
- Subset 3 (15 taxa): *Cunninghamia lanceolata, Metasequoia glyptostroboides, Sequoiadendron giganteum, Taxodium distichum, Juniperus scopulorum, Calocedrus decurrens, Platycladus orientalis, Thuja plicata, Austrocedrus chilensis, Pilgerodendron uviferum, Callitris macleayana, Callitris gracilis, Taiwania cryptomerioides, Trebouxia arboricola, Stephanosphaera pluvialis*
- Subset 4 (16 taxa): *Athrotaxis cupressoides, Sequoia sempervirens, Cryptomeria japonica, Glyptostrobus pensilis, Tetraclinis sp., Microbiota decussata, Cupressus dupreziana, Fokienia hodginsii, Thujopsis dolabrata, Papuacedrus papuana, Neocallitropsis pancheri, Diselma archeri, Widdringtonia cedarbergensis, Chamaecyparis lawsoniana, Leptosira obovata, Brachiomonas submarina*
- Subset 5 (19 taxa): *Ginkgo biloba, Encephalartos barteri, Stangeria eriopus, Picea engelmannii, Abies lasiocarpa, Keteleeria evelyniana, Cystopteris reevesiana, Huperzia squarrosa, Selaginella wallacei, Pinus radiata, Pinus taeda, Amborella trichopoda, Nuphar advena, Nymphaea sp., Pseudotsuga wilsoniana, Ephedra sinica, Welwitschia mirabilis, Microthamnion kuetzingianum, Haematococcus pluvialis*
- Subset 6 (20 taxa): *Cycas micholitzii, Dioon edule, Cedrus libani, Pseudolarix amabilis, Nothotsuga longibracteata, Tsuga heterophylla, Nephrolepis exaltata, Dennstaedtia davallioides, Selaginella moellendorffii, Pinus parviflora, Pinus ponderosa, Pinus jeffreyi, Austrobaileya scandens, Kadsura heteroclita, Illicium floridanum, Cathaya argyrophylla, Larix speciosa, Gnetum montanum, Botryococcus terribilis, Chlamydomonas noctigama*

**Fig. S6.1.**
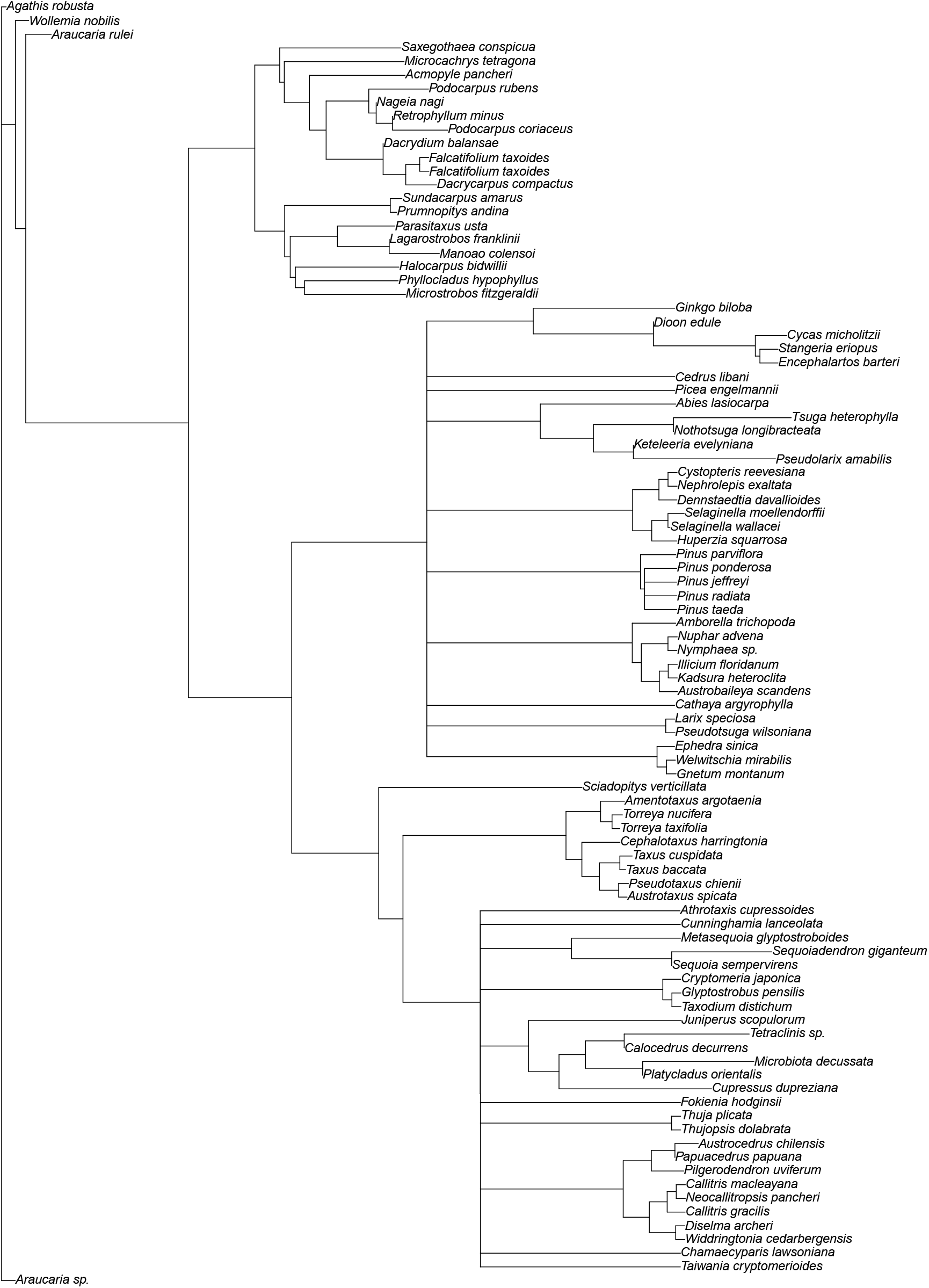
Gymnosperm tree of blobs inferred by TINNiK (*α* = 0.01, *β* = 0.99).

**Fig. S6.2.**
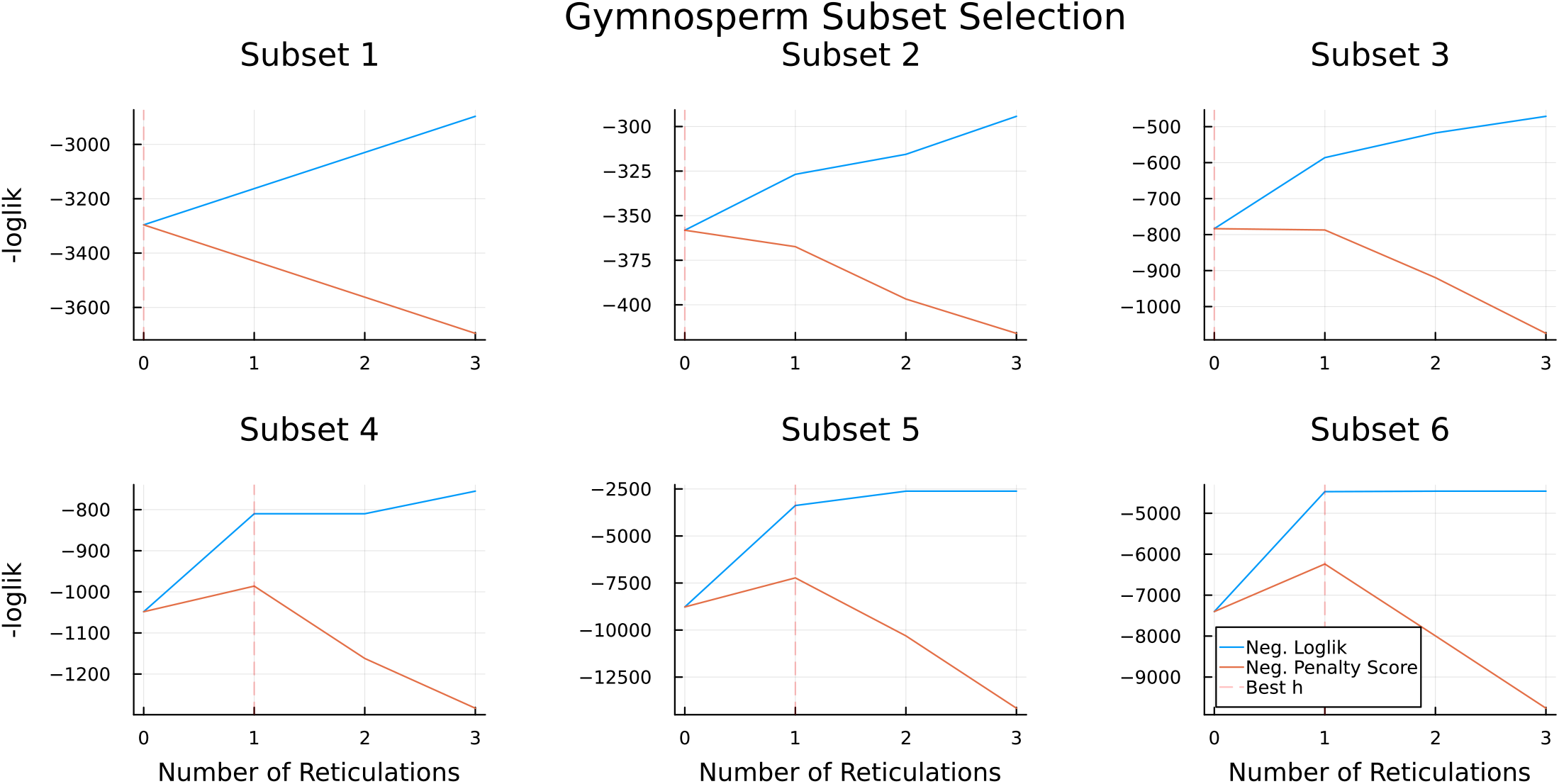
Model selection figures corresponding to each of the 6 subsets listed above. The blue line is the negative log pseudolikelihood of each model (higher is better), the orange line is the penalized score for each model (higher is better), and the vertical dashed red line is placed at the value of *h* with the largest penalized score. This is the model that was selected.

#### Ferns

- Subset 1 (22 taxa): *Dendrolycopodium obscurum, Pseudolycopodiella caroliniana, Huperzia selago, Huperzia lucidula, Huperzia squarrosa, Chloromonas tughillensis, Oogamochlamys gigantea, Volvox aureus, Vitreochlamys sp., Pleurastrum insigne, Welwitschia mirabilis, Cupressus dupreziana, Thujopsis dolabrata, Diselma archeri, Falcatifolium taxoides, Dacrydium balansae, Halocarpus bidwillii, Prumnopitys andina, Pinus jeffreyi, Pseudolarix amabilis, Ettlia oleoabundans, Dunaliella tertiolecta*
- Subset 2 (7 taxa): *Selaginella moellendorffii, Selaginella willdenowii, Selaginella cf. pallescens, Isoetes sp., Isoetes tegetiformans, Parachlorella kessleri, Dunaliella primolecta*
- Subset 3 (23 taxa): *Osmunda javanica, Osmunda sp., Osmundastrum cinnamomeum, Marattia attenuata, Angiopteris evecta, Danaea nodosa, Dipteris conjugata, Hymenophyllum bivalve, Ophioglossum petiolatum, Sceptridium dissectum, Botrypus virginianus, Psilotum nudum, Tmesipteris parva, Lygodium japonicum, Anemia tomentosa, Pilularia globulifera, Azolla cf. caroliniana, Dennstaedtia davallioides, Didymochlaena truncatula, Leucostegia immersa, Nephrolepis exaltata, Prototheca wickerhamii, Dunaliella salina*
- Subset 4 (23 taxa): *Alsophila pinulosa, Plagiogyria japonica, Culcita macrocarpa, Thyrsopteris elegans, Lonchitis hirsuta, Lindsaea microphylla, Lindsaea linearis, Cryptogramma acrostichoides, Argyrochosma nivea, Myriopteris rufa, Notholaena montieliae, Parahemionitis cordata, Gaga arizonica, Vittaria lineata, Vittaria appalachiana, Adiantum aleuticum, Adiantum raddianum, Ceratopteris thalictroides, Pityrogramma trifoliata, Pteris vittata, Pteris ensiformis, Picochlorum atomus, Dunaliella salina*
- Subset 5 (12 taxa): *Davallia fejeensis, Phymatosorus grossus, Pleopeltis polypodioides, Phlebodium pseudoaureum, Polypodium amorphum, Polypodium hesperium, Polypodium hesperium, Polypodium glycyrrhiza, Bolbitis repanda, Polystichum acrostichoides, Eremosphaera viridis, Dunaliella salina*
- Subset 6 (21 taxa): *Homalosorus pycnocarpos, Thelypteris acuminata, Deparia lobato-crenata, Diplazium wichurae, Blechnum spicant, Athyrium filix-femina, Athyrium sp., Onoclea sensibilis, Woodsia ilvensis, Woodsia scopulina, Asplenium platyneuron, Asplenium nidus, Asplenium cf. x lucrosum, Cystopteris protrusa, Cystopteris fragilis, Cystopteris reevesiana, Cystopteris fragilis, Cystopteris utahensis, Gymnocarpium dryopteris, Nannochloris atomus, Asteromonas gracilis*

**Fig. S6.3.**
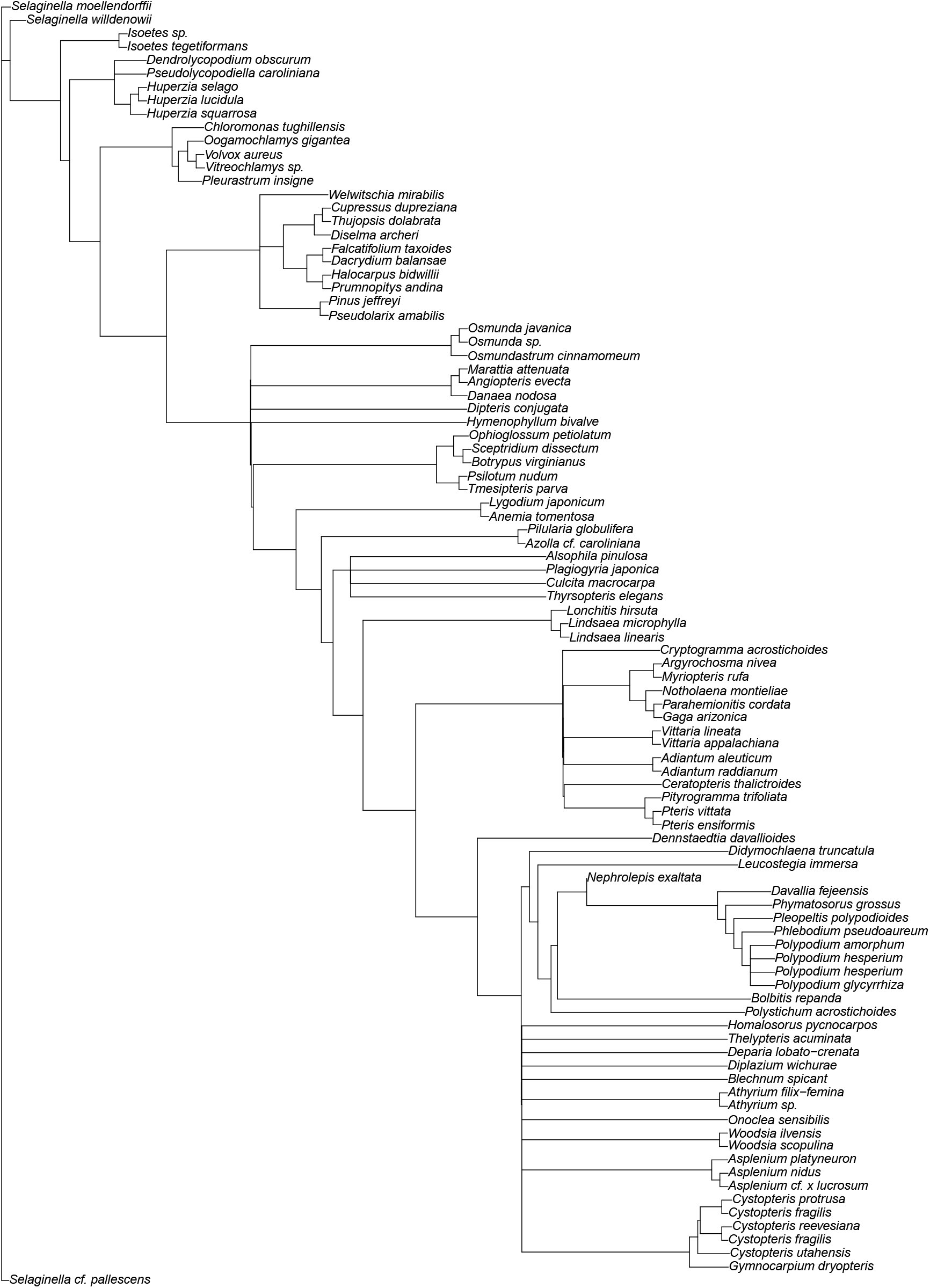
Fern tree of blobs inferred by TINNiK (*α* = 0.001, *β* = 0.999).

**Fig. S6.4.**
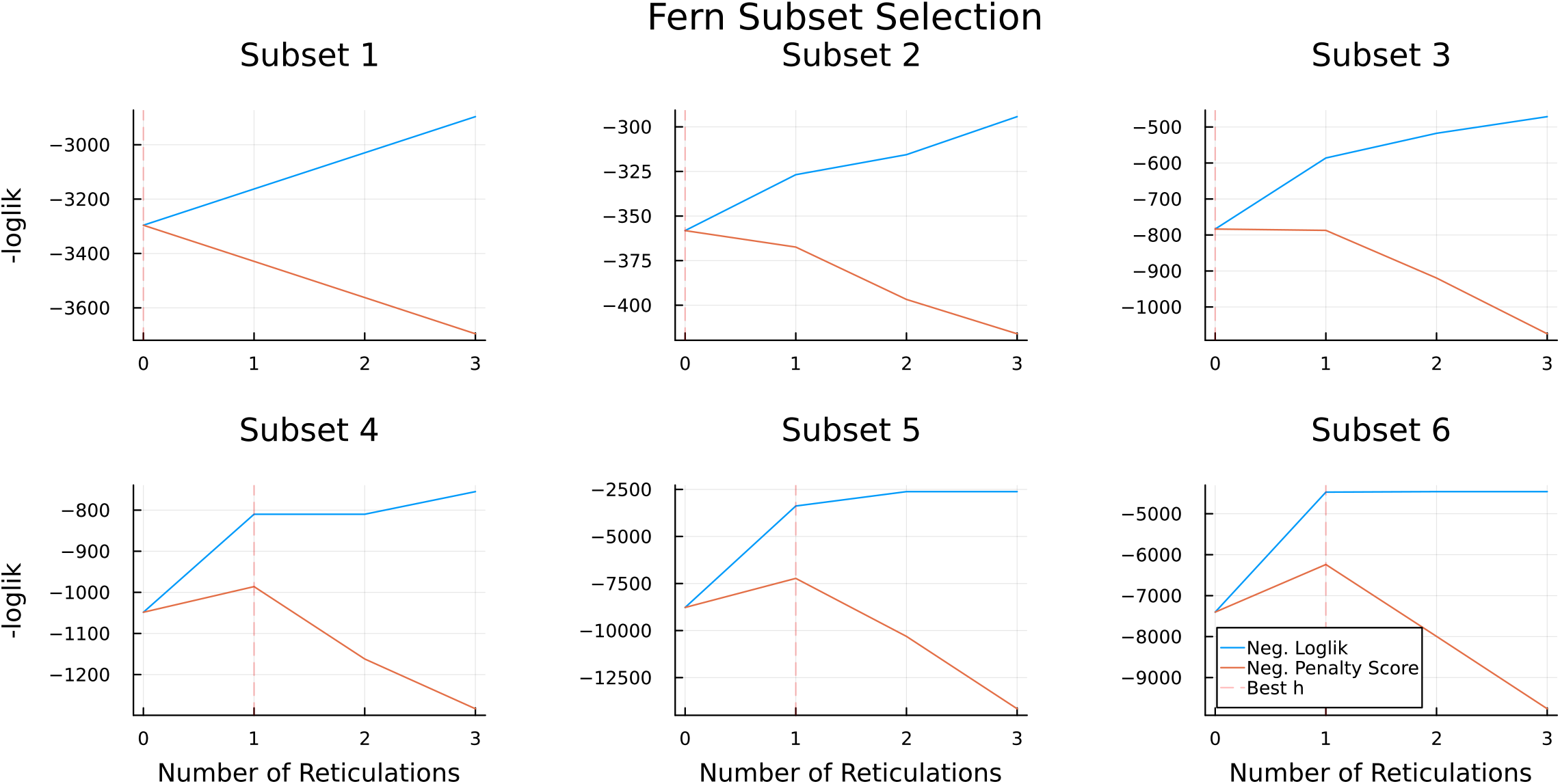
Model selection figures corresponding to each of the 6 subsets listed above. The blue line is the negative log pseudolikelihood of each model (higher is better), the orange line is the penalized score for each model (higher is better), and the vertical dashed red line is placed at the value of *h* with the largest penalized score. This is the model that was selected.

